# The Bgee suite: integrated curated expression atlas and comparative transcriptomics in animals

**DOI:** 10.1101/2020.05.28.119560

**Authors:** Frederic B. Bastian, Julien Roux, Anne Niknejad, Aurélie Comte, Sara S. Fonseca Costa, Tarcisio Mendes de Farias, Sébastien Moretti, Gilles Parmentier, Valentine Rech de Laval, Marta Rosikiewicz, Julien Wollbrett, Amina Echchiki, Angélique Escoriza, Walid H Gharib, Mar Gonzales-Porta, Yohan Jarosz, Balazs Laurenczy, Philippe Moret, Emilie Person, Patrick Roelli, Komal Sanjeev, Mathieu Seppey, Marc Robinson-Rechavi

## Abstract

Bgee is a database to retrieve and compare gene expression patterns in multiple animal species, produced by integrating multiple data types (RNA-Seq, Affymetrix, in situ hybridization, and EST data). It is based exclusively on curated healthy wild-type expression data (e.g., no gene knock-out, no treatment, no disease), to provide a comparable reference of normal gene expression. Curation includes very large datasets such as GTEx (re-annotation of samples as “healthy” or not) as well as many small ones. Data are integrated and made comparable between species thanks to consistent data annotation and processing, and to calls of presence/absence of expression, along with expression scores. As a result, Bgee is capable of detecting the conditions of expression of any single gene, accommodating any data type and species. Bgee provides several tools for analyses, allowing, e.g., automated comparisons of gene expression patterns within and between species, retrieval of the prefered conditions of expression of any gene, or enrichment analyses of conditions with expression of sets of genes. Bgee release 14.1 includes 29 animal species, and is available at https://bgee.org/ and through its Bioconductor R package BgeeDB.

## INTRODUCTION

Gene expression is central to the relation between genes or genomes and their function. It mediates or indicates the involvement of genes in functions, pathways, pathologies, differences between individuals, species, or genes. Many of the features which make expression so interesting also complicate its bioinformatics analysis, especially for multicellular organisms such as animals. While databases of reference genomes, providing standardized annotations and comparative frameworks within and between species, are well established(1–3), this is not the case for expression data. Both the concept of a “reference expression” for a species, and comparisons, are complicated by the need to define comparable samples and conditions(4).

There are a few datasets which can play a role close to that of a reference expression pattern. Such “atlases” cover typically many anatomical structures (tissues or organs), and are often used to present expression in gene-centric databases, such as NCBI Gene(5). Gene presents by default expression from the HPA set(6) for human genes, with the option of switching to another of three RNA-seq atlases, covering 6 to 27 tissues. Because they are presented separately, it can be difficult to come to a conclusion concerning the expression pattern of a gene. For example, human insulin (NCBI Gene ID: 3630) shows very pancreas-specific expression from HPA, but the other proposed atlases lack pancreas samples, and show top expression in the prostate, the spleen, or the stomach. Probably the most complete atlas so far is GTEx(7). While atlases are valuable, they are dispersed in different databases, do not have common standards of annotation or quality-control, provide differently processed values, and cover different limited subsets of anatomy and conditions.

There exist databases which specialize in providing expression pattern descriptions(8). The Expression Atlas(9) presents curated and processed expression data from a diversity of species and sources. As of May 2020, it includes 3,942 studies from 65 species, of which 214 are annotated as “baseline”, and are used to present an expression pattern of each gene. Expression is presented per experiment. Tissues(10) presents expression patterns derived from UniProtKB annotations, selected atlases, and text mining. The focus is on confidence in expression presence rather than levels of expression. Data from different techniques is presented separately. An interesting feature is that tissues are ranked within each evidence type, allowing the most important features of a complex expression pattern to emerge. Thus, for human insulin, the pancreas is one of two tissues from UniProtKB and Pancreatic beta cell is the top tissue from text mining. Finally, Model Organism Databases (MODs)(11)(12) provide integration of many datasets for a given species (or a targeted set of species). They integrate many small scale experiments such as in situ hybridization, thanks to curation and the pioneering use of ontologies. They do not cover species outside of a small core of model organisms, and include expression data both from mutant strains and from wild-type.

We have developed the Bgee database to answer questions about gene expression while avoiding limitations due to the separation of data sets and species. We notably provide an integrated reference of gene expression patterns in animals. We capture conditions of gene expression, i.e., the biological parameters for which a gene is expressed. Conditions are combinations of sample features, which together are expected to define an expression state. As of Bgee 14, we capture conditions as unique combinations of species, anatomical structure, developmental stage, sex and strain (sex and strain are not yet accessible to users in all views or tools). The data produced by Bgee allow: i) to answer the questions “where and when this gene is expressed?”, and “what are the genes expressed in this condition?”; ii) to identify the conditions most relevant to the expression of each gene; iii) to study gene expression evolution, by comparing expression patterns between species. Bgee provides a consistent vision over all species, and integrates data over a large number of atlases and small-scale datasets. These currently include 29 species and four major types of expression experimental data.

## MATERIALS AND METHODS: DATA CURATION AND PROCESSING

### Pipeline overview

We collect expression data from different sources, depending on their data type. Hereafter we define “data type” to cover data resulting from different types of assays for gene expression. We curate these data to retain only samples from healthy wild-type individuals. We annotate them to anatomical and life stage ontologies, along with information of population/strain and sex. We perform quality controls to remove low-quality and duplicated samples. We process these data to produce present/absent expression calls, along with expression level information, represented by our expression ranks and expression scores. We propagate these calls along the anatomical and life stage ontologies, to allow the integration of data generated with various granularities. We provide an overview of this pipeline in Figure 1, and detail some aspects in the following subsections; see the Supplementary Methods and the online documentation for further details.

**Figure 1:**
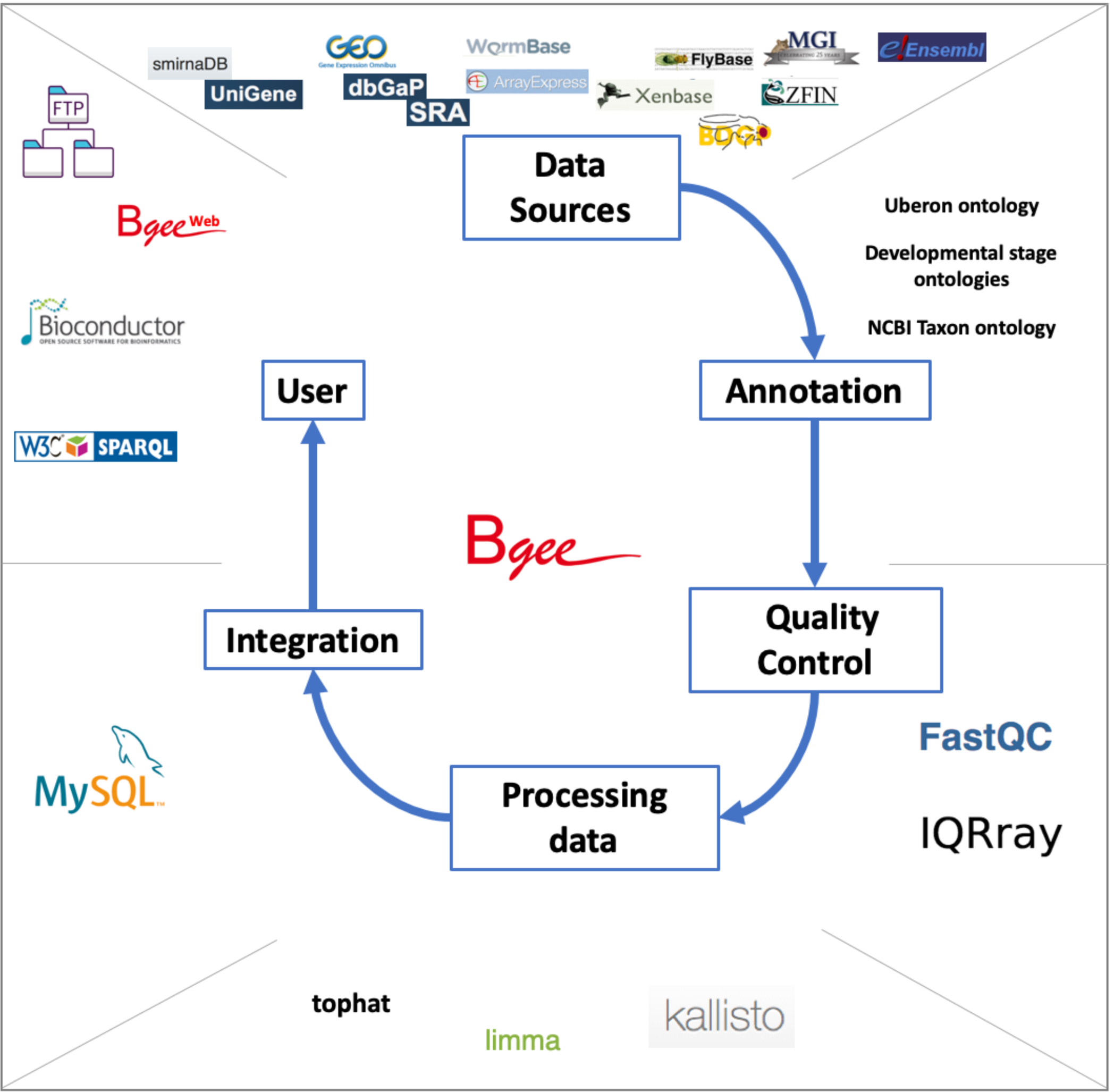
Bgee pipeline overview. Expression data are retrieved from various databases; they are annotated by the Bgee team, or annotations from Model Organism Databases are remapped by the Bgee team, to ontologies describing developmental stages, anatomy, taxa; quality controls are performed, using for instance FastQC for RNA-Seq data, IQRray for Affymetrix data; data are then analyzed using specific tools, such as kallisto to produce TPM values from RNA-Seq data, or limma to compute TMM normalization factors, and presence/absence expression calls are then produced; all the expression data and analysis results are integrated into the MySQL Bgee database; these data are then leverated by the different tools offered by Bgee: Bgee web-interface, Bioconductor packages, SPARQL endpoint, FTP server. Icons for tools and databases retrieved from their respective website.

Ontologies and genome annotations are only updated for each major release (e.g., Bgee 13 to 14), while expression data can be updated at minor releases (e.g. Bgee 14.0 to 14.1). Thus all expression data presented here concern Bgee 14.1 specifically, while ontology and genome annotations concern Bgee 14, both 14.0 and 14.1.

### Species information: taxonomy, genomes, gene orthology

We annotate species information using the NCBI taxonomy database(13), and integrate the taxonomy for species present in Bgee. We retrieve gene models for each species in Bgee from Ensembl(3) (Ensembl 84 for Bgee 14) and Ensembl Metazoa(2) (release 30 for Bgee 14) using the Perl Ensembl API. We also retrieve cross-references to other databases, gene biotypes, and annotations to Gene Ontology(14)(15) terms, and gene orthology information from OMA(16), although our web and R views and tools do not make use of GO annotations nor of orthology as of Bgee 14.1.

### Uberon integration

To describe the anatomy of diverse animals, we use the Uberon ontology(17). Its integration into Bgee requires processing, described in Supplementary Methods. Briefly, we refine taxon constraints; we simplify Uberon to keep only the knowledge needed for Bgee, and simplify it for an easier browsing; we remove any cycles; and we correct mappings between model organism databases (MODs) and Uberon.

### Developmental stage integration

Each species in Bgee has a developmental stage ontology, available in the obophenotype developmental stage ontology repository. We have developed all of the species-specific ontologies in this repository, except those from Model Organism Databases (EMAPA for mouse(18); FBdv for fly(19); ZFA for zebrafish(20); XAO for Xenopus(21, 22); WBls(23) for worm). We produce a composite stage ontology merging all of these species-specific ontologies, that we can insert into Bgee. Having one common multi-species ontology, with high level terms merged between species (e.g., “gastrula”), allows us to propagate expression data to comparable stages in different species, and thus take development into account when automatically comparing expression patterns. The merge is performed by the Uberon maintainers.

We describe our strategy for insertion into Bgee in detail in the Supplementary Methods. Briefly, we merge terms from species-specific ontologies into the structure of Uberon; we order stages using the relations “preceded_by” or “immediately_preceded_by”; and we ensure that each stage has at most one parent by a *part_of* relation. Finally, we transform the merged developmental stage ontology into a nested set model for integration into Bgee.

### Curation of expression data

Our curation steps have the aim of selecting data: i) produced from healthy wild-type animals in normal conditions; ii) that are of high quality; iii) that are non-redundant. Because the amount of data available in public repositories is large, we can be stringent regarding our criteria to accept data for inclusion.

#### Data sources

All data sources are detailed in Table 1. For SRA or GEO studies for which the corresponding publication is missing at time of curation by Bgee, we determine the publication, and report missing PMIDs to GEO. We use the information from these publications, as well as those which are already linked, for precise annotation of samples to anatomy, stage, sex and/or strain. For example the SRP075519 study contains six RNA-seq libraries, with tissue information reported as ‘muscle’, while the related paper(24) reports that “RNA-Seq analysis was performed on six piglets representing two breeds: Duroc and Ronghcang (three animals of each breed). RNA was extracted from the longissimus dorsi of each individual.” We can thus annotate the libraries to the relevant term UBERON:0001401 “longissimus thoracis muscle”, and we can also annotate each library to the corresponding breed (here ‘Duroc’ or ‘Ronghcang’).

**Table 1:**
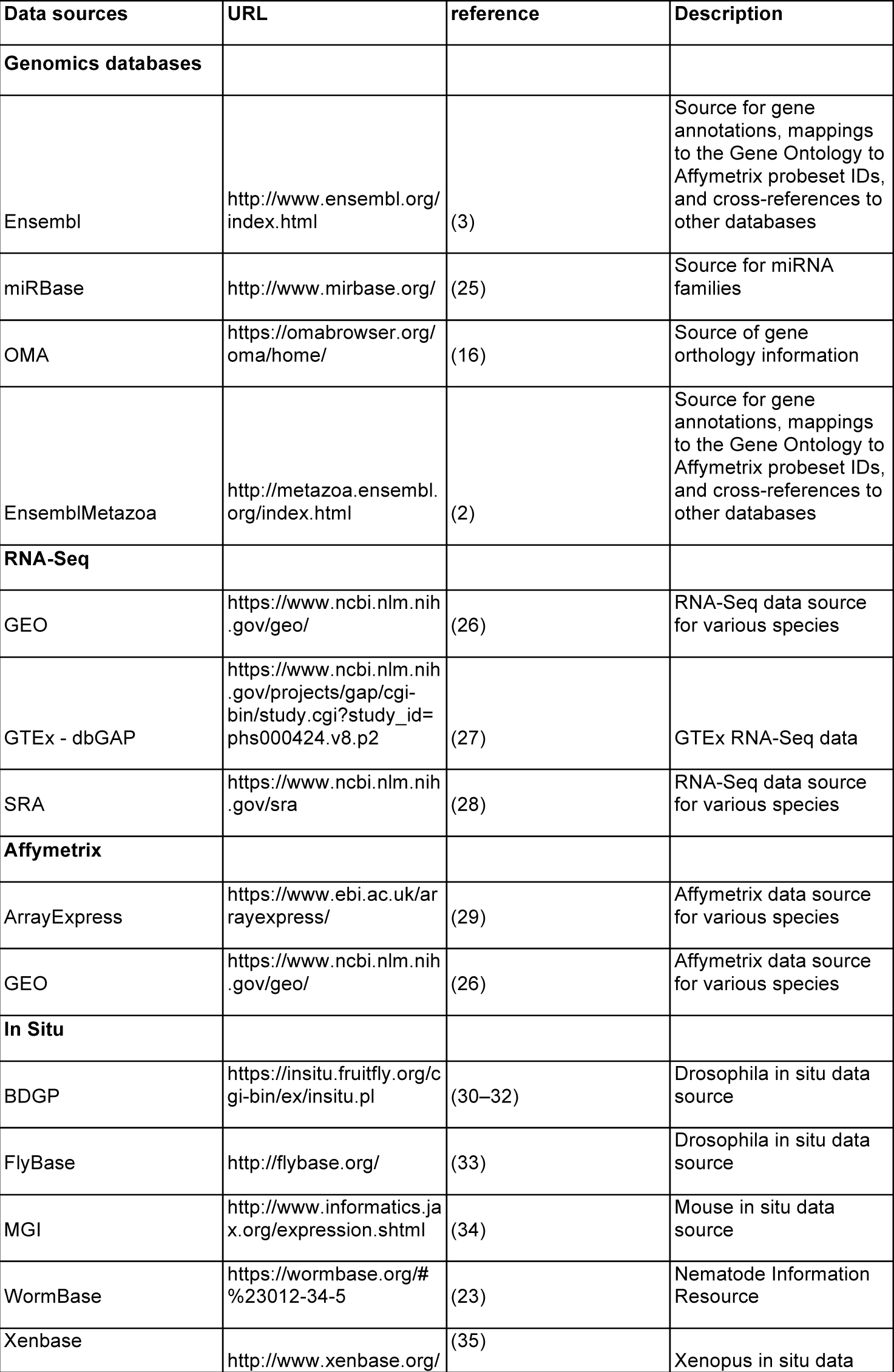
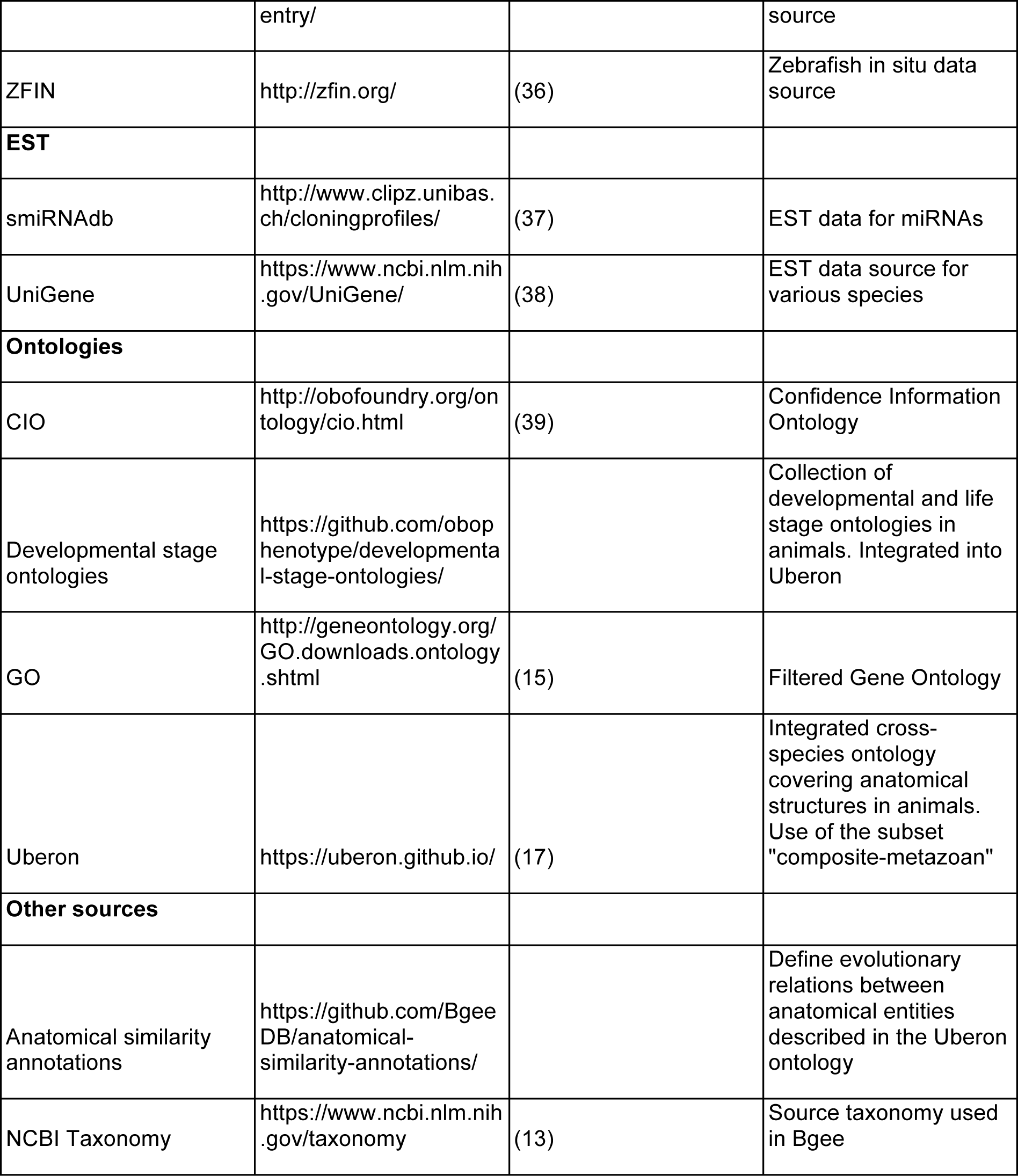
Data sources of Bgee, with URLs, references, and descriptions.

#### Healthy wild-type samples in normal conditions

For inclusion into Bgee, we reject samples from animals with abnormal genetic backgrounds (e.g., project SRX2751118, nono^-/-^ mice), or subject to diseases, to gene knockouts, or to treatments not expected in the wild (e.g., irradiated sample DRX012402). We manually review the information about each sample before inclusion (see Supplementary Material for our criteria for inclusion). While we aim at being stringent regarding the quality of the data we integrate, we also need our data to reflect the genetic diversity of wild-type animals, or the various expected conditions encountered in the wild.

For *in situ* hybridization data that we retrieve from MODs, we coordinate with developers of these resources to find the appropriate parameters to retrieve only wild-type healthy samples in normal conditions.

#### Removal of hidden redundancies

During our quality control procedure, we have identified duplicated content in the GEO and ArrayExpress databases (identical arrays reused in different experiments, and submitted as distinct), affecting about 14% of our Affymetrix data(40). We have thus created a method to discard these duplicated samples, as well as to avoid this problem appearing in our RNA-Seq dataset. This allows Bgee, despite the use of potentially overlapping multiple datasets, to provide and use a clean reference of unique samples for downstream analyses.

#### Curation of GTEx dataset

We have curated the GTEx human dataset phs000424.v6.p1(7). We apply a stringent re-annotation process to the GTEx data to retain only healthy and non-contaminated samples, using the information available under restricted-access. For instance, we reject all samples for 31% of subjects, deemed globally unhealthy from the pathology report (e.g., drug abuse, diabetes, BMI > 35), as well as specific samples from another 28% of subjects who had local pathologies (e.g., brain from Alzheimer patients). We also reject samples with contamination from other tissues according to the pathologist report. In total, we have kept only 6008 of 11983 samples (50%); these represent a high quality subset of GTEx, restricted to only healthy and non-contaminated samples. We re-annotate some GTEx samples to Uberon ontology, according to the original sampling sites. For instance, we map GTEx samples ‘Minor Salivary Gland - Inner surface of lower lip’ to UBERON:0001830 minor salivary gland, while GTEx reports on UBERON:0006330 anterior lingual gland. Note that both lip inner surface and tongue have minor salivary glands. The precise criteria for our curation of GTEx can be found in Sup. Material. More information to use these data is available at https://bgee.org/?page=doc&action=data_sets, and is also provided in Sup. Material.

### Annotation of expression data

We capture information about the anatomical localization of samples, their developmental and life stage, their sex, and their strain or ethnicity. We either manually capture this information using ontologies and controlled vocabularies (for Affymetrix, RNA-Seq, and EST data), or we map existing annotations provided by MODs to these ontologies and vocabularies (for all *in situ* hybridization data, and for some Affymetrix and RNA-Seq data annotations provided by Wormbase for *C. elegans*). For Affymetrix and RNA-Seq data, we also capture whether replicates are technical or biological.

In addition to Uberon^22^ and developmental stage ontologies described above, sex is annotated to a simple controlled vocabulary, and we maintain a naive controlled vocabulary of strains or breeds, based on the UniProt ‘strains.txt’ file (https://www.uniprot.org/docs/strains) and completed with strain/breed names found in the literature, or from species specific resources (e.g. https://www.gov.uk/guidance/official-cattle-breeds-and-codes). We standardize strain information provided in free-text format, notably to remove most duplicates, resulting in 422 different strain and ethnicity terms used in Bgee 14.1. If strain or sex information are missing, they are mapped to one of the following: “NA”, not available from source information; “not annotated”, information not captured by Bgee; “confidential_restricted_data”, information cannot be disclosed publicly (e.g., GTEx).

### Processing of expression data

We present key processing steps and software very briefly here. Details of expression data download and processing are in Supplementary Methods, and in the online code documentation. For RNA-seq, we use Ensembl and Ensembl metazoa for sequences and annotations of coding and non coding genes. We use Kallisto(41) to generate pseudo-counts per transcript, which are then summed for each gene. All further analysis and reporting is done at the gene level as of Bgee 14. For Affymetrix microarray data, we use CEL files when they are available, using IQRray(42) for quality control and gcRMA(43) for normalization. Otherwise we use MAS5(44) processed files. Mapping of probesets to genes is recovered from Ensembl. For ESTs, we simply mapped by BLAST(45) to UniGene clusters then to Ensembl genes, and counted the number of ESTs per gene.

### Integration of *in situ* hybridization data from Model Organism Databases

We retrieve *in situ* hybridization data from the relevant MODs (Table 1), using InterMine(46) whenever possible. We filter out the data from animals with abnormal genetic backgrounds, e.g., animals with transgenes or knockout. We remap species-specific terms from the MOD ontology or controlled vocabulary to Uberon. When needed, we transform annotations to separate anatomical localization and developmental stage. We provide details of this integration, and of specific issues and their solutions, in Supplementary Methods.

### Calls of presence/absence of expression

Bgee provides calls of presence or absence of expression for unique combinations of a gene in a condition. As of Bgee 14.1, a condition is defined by 5 pieces of information: an anatomical entity, a life stage, a sex, a strain or ethnicity, and a species.

#### Calls produced from RNA-Seq data

We use a new method to estimate for each RNA-Seq library independently the TPM threshold to consider a gene as actively transcribed (Julien Roux, Marta Rosikiewicz, Julien Wollbrett, Sara S. Fonseca Costa, Marc Robinson-Rechavi, Frederic B. Bastian; in preparation). While this method will be described elsewhere, the documentation of our pipeline describing its use in Bgee is available at https://github.com/BgeeDB/bgee_pipeline/tree/master/pipeline/RNA_Seq (archived version for release Bgee 14.1: https://github.com/BgeeDB/bgee_pipeline/tree/v14.1/pipeline/RNA_Seq). Briefly, we use all RNA-seq data from a species to identify a stringent set of intergenic regions, then use these regions to define the background level of read mapping per library. Then, we call genes as expressed when their level of mapped reads is significantly higher than this background, and as not expressed otherwise. As of Bgee 14.1, RNA-Seq calls are all considered of high quality.

#### Calls produced from Affymetrix data

When only the MAS5 files of an analysis are available, we use the flags provided by the MAS5 software(47). Although MAS5 classification is efficient(48), the estimation of the background signal can be biased depending on probe sequence affinity(49). For this reason, we use preferentially CEL files when available to produce present/absent calls, and, when not available, we consider all calls produced from MAS5 files as low quality. MAS5 “present” and “marginal” flags correspond in Bgee to a low quality present call; “absent” flag to a low quality absent call. For CEL data, we use the gcRMA algorithm(43) to normalize the signal taking into account probe sequences, and use a subset of weakly expressed probesets for estimating the background, as described in (49). We then apply a Wilcoxon test to compare the normalized signal of the probesets with the background signal, as implemented in the ‘mas5calls’ function of the bioconductor package ‘affy’(48): High quality present call corresponds to a p-value threshold of < 0.03 (corresponding to a FDR of ≈5%, see Schuster *et al*.(49)), low quality present call to a p-value >= 0.03 and <= 0.12, and high quality absent call to a p-value > 0.12 (< 0.12 corresponds to a FDR ≈ 10%).

We then exclude all probesets that are never seen as MAS5 “present” or CEL high quality present over the whole dataset. Because a same gene can be covered by several probesets, we reconcile this information to produce one call per gene and per chip, by retaining the best probeset signal, in this order: high quality present, low quality present, high quality absent, low quality absent.

#### Calls produced from EST data

Based on the number of ESTs mapped to a gene for which the 95% confidence interval of the EST count excludes 0 (50), we call presence of expression of high quality when an experiment has at least 7 ESTs mapped to a gene, and of low quality from 1 to 6 ESTs. We do not produce calls of absence of expression from EST data because of the low sampling.

#### Remapping to more generalized conditions

We annotate and retrieve expression data with as much granularity as possible. But such a granularity can hamper call integration and comparison. Thus, we remap the granular annotations of the data to more generalized conditions. For developmental stages, we have created in developmental stage ontologies a subset named “granular_stage”. For instance, HsapDv:0000150 “56-year-old human stage” would be automatically remapped to HsapDv:0000092 “human middle aged stage”. The highest granularity possible is still available when retrieving our annotated and processed expression values.

#### Call propagation

After producing these calls from multiple techniques and experiments, we propagate them along the graph of conditions (see Figure 2 for an overview). We produce the graph of conditions by: i) using the graph of anatomical entities (Uberon ontology) from *is_a* (subClassOf in OWL) and *part_of* relations (an object property in OWL); ii) using the graph of developmental stages (developmental stage ontology) from *part_of* relations; iii) adding a root parent to all sex terms by *is_a* relation; iv) adding a root “wild type” parent to all strain terms by *is_a* relation (in the context of Bgee, all strains have Wild-Type as parent).

**Figure 2:**
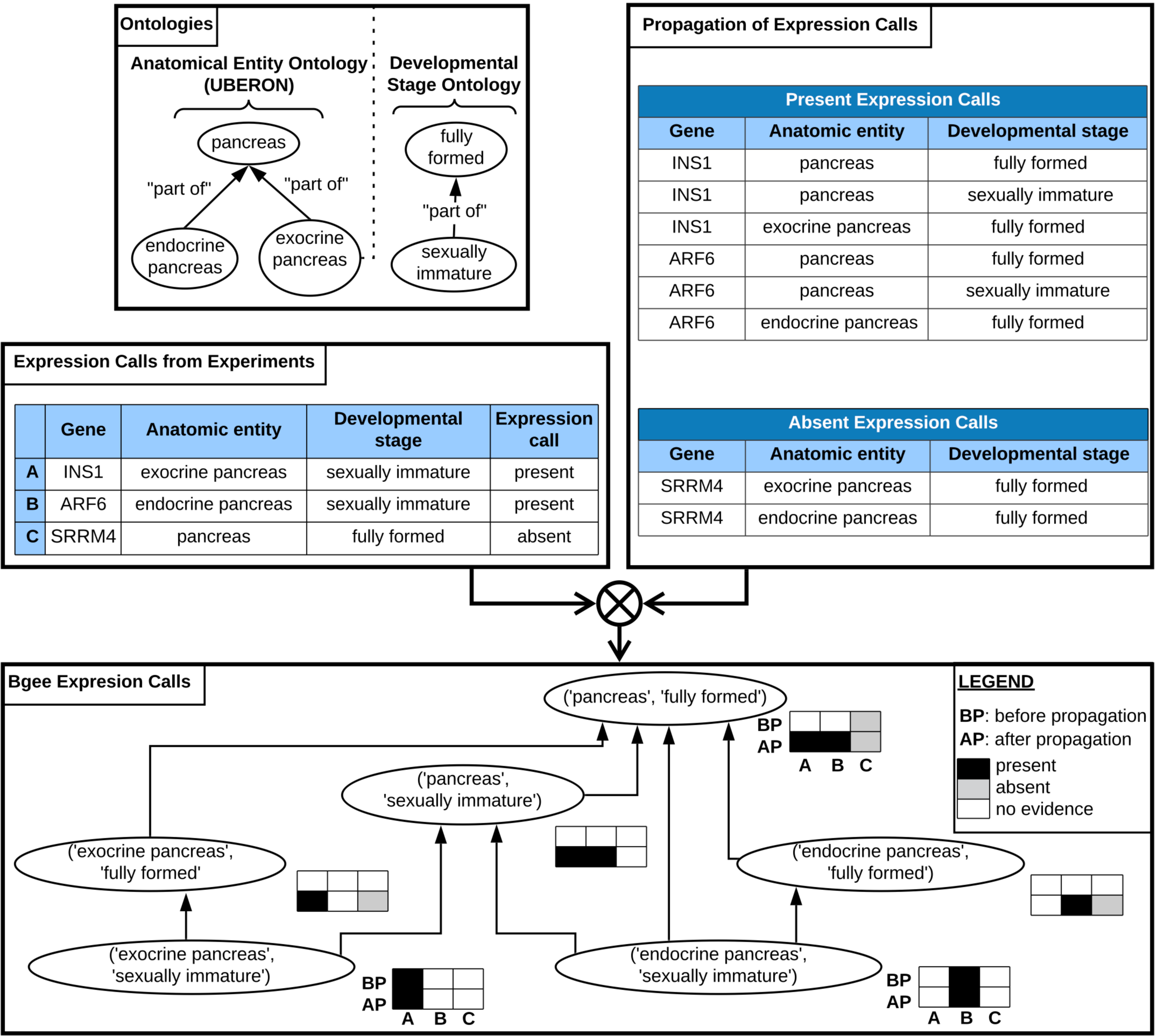
propagation of calls of presence/absence of expression. Calls of presence/absence of expression are produced from the raw data (left table), for instance: call of presence of expression for gene INS1 in exocrine pancreas at sexually immature developmental stage; call of presence of expression for gene ARF6 in endocrine pancreas at sexually immature developmental stage; call of absence of expression for gene SRRM4 in pancreas at fully formed developmental stage. A graph of conditions is generated by using the anatomical ontology and the developmental stage ontology, to allow propagation of expression calls (top left box): for instance, the condition “endocrine pancreas (UBERON:0000016) - sexually immature (UBERON:0000112)” is a child of the condition “pancreas (UBERON:0001264) - fully formed (UBERON:0000066)”; the condition “endocrine pancreas (UBERON:0000016) - fully formed (UBERON:0000066)” is a parent of the condition “endocrine pancreas (UBERON:0000016) - sexually immature (UBERON:0000112)”. Calls of presence of expression are propagated to all parent conditions; calls of absence of expression are propagated to direct child anatomical entities (top right box). The bottom box shows the hierarchy of conditions, and how data are propagated. This propagation of calls allow the integration of data that were produced and annotated with different granularity: for instance, while before propagation there was information in “pancreas (UBERON:0001264) - fully formed (UBERON:0000066)” only for the gene SRRM4, after propagation the expression of the three genes can be compared in this condition (bottom box).

We propagate calls of presence of expression along the graph of conditions to all parent conditions. The idea is that if a gene is expressed, e.g., in the midbrain, it is expressed in the brain, the parent structure. We propagate calls of absence of expression to direct anatomical sub-structures, keeping the same developmental stage, sex, and strain. We propagate only to direct anatomical sub-structures, and not all sub-structures, because we consider that an experiment could miss the expression of a gene in a very small part of a larger anatomical entity. We do not propagate to child terms for stages, sexes, or strains.

#### Integration to produce one global call and confidence level per gene - condition combination

Multiple calls from multiple data types and assays can be produced for a given gene in a given condition, from direct observation or from propagation along the condition graph. We integrate all these individual calls to produce one global call of presence/absence of expression per gene - condition, along with a confidence level.

Global calls of absence of expression are reported when all experiments consistently report an absence of expression for a gene in a condition, with no conflicting presence of expression reported in the condition itself, or any sub-condition. Otherwise, a global call of presence of expression is reported: presence of expression always “wins” over absence of expression, whatever the quality level and number of experiments producing the call of presence of expression, as of Bgee 14. This means that Bgee is very stringent when reporting absence of expression.

We follow the principles of the Confidence Information Ontology (CIO)(39), and translate them into three confidence levels. After call propagation and reconciliation of presence/absence calls, we assign gold confidence level to global calls supported by at least two different experiments with a high quality call; silver to global calls supported by either only one experiment with a high quality call, or at least two different experiments with a low quality call; and bronze level to global calls supported by only one experiment with a low quality call. Because of call propagation, we also consider in support of a global call the experiments which contribute to calls in sub-conditions (for calls of presence of expression) or in direct parent anatomical entities (for calls of absence of expression).

### Expression ranks and expression scores

We also provide a quantitative ranking of the conditions where a gene is expressed, also integrated over experiments and data types. Briefly, these ranks are computed in each condition relative to other genes, which then allows to compare conditions for each gene (after normalization). The assumption is that if a gene has a lower rank (i.e., higher relative expression) in some conditions than in others, then its expression is more important in these conditions. To compute expression ranks: 1) we compute ranks using different methods per data type; 2) we normalize ranks over all genes, conditions and data types for each species; 3) we compute a global weighted mean rank for each gene in each condition over all data types considered. We also transform these ranks into expression scores, more easily understandable by users: higher gene expression translates into lower rank but higher expression score, from 0 to 100.

We present briefly the principle of our rank computation. Detailed methods are available in Supplementary Methods and online code documentation.

#### Rank computation for each data type

For RNA-Seq, we compute fractional ranks of genes in each library, based on their TPM value. Then for each gene and each condition with RNA-Seq data in the condition itself, we compute a weighted mean of the gene ranks over relevant libraries. We weigh the mean by the number of distinct ranks in each library, under the assumption that libraries with a higher number of distinct ranks have a higher power for ranking genes (see eq. 1).

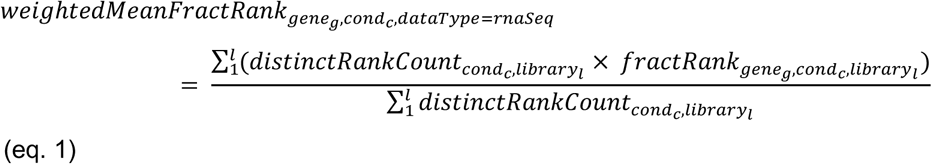

where *distinctRankCount* is the count of distinct ranks in a library, and *fractRank* the fractional rank of a gene in a library.

Similarly, for Affymetrix we compute fractional ranks for each chip based on signal intensity, considering the highest signal intensity of the probesets for each gene. We normalize the fractional ranks between chip types, to correct for different genomic coverages due to different probeset designs. First, for each chip type, we compute its max rank over all conditions. Then, we normalize ranks of each chip, based on the max rank of this chip type, as compared to the max of the max ranks of other chip types represented in the same condition (see eq. 2; note that we do not normalize based on the max rank in a given condition, but on the max rank of the chip types represented in the condition). We then compute the weighted mean of the normalized ranks per gene and condition, weighting by the number of distinct ranks in each chip (see eq. 3).

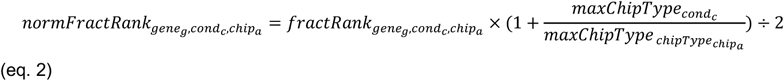

where *fractrank* is the fractional rank of a gene in a chip,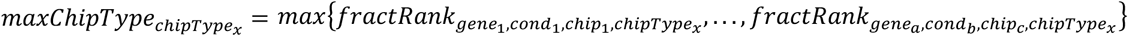(the max of the chip type over all conditions) and 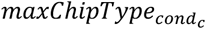is 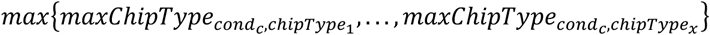 (the max of the chip types represented in the condition)

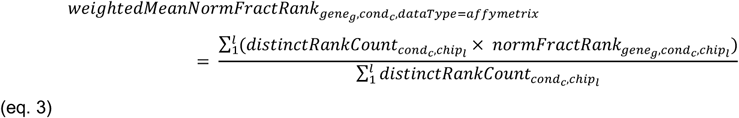

Since *in situ* hybridization data are not quantitative, we make the assumption that the more often an expression observation is reported, the more biologically important this expression is likely to be. Thus the rank is computed on the number of calls and their confidence pulling together all data available in each condition, using a dense ranking instead of a fractional ranking. We thus directly obtain a rank for each gene and condition, without averaging values as for RNA-Seq and Affymetrix.

While EST data are in principle quantitative, their very low coverage leads us to treat them similarly to *in situ* hybridization data, summing EST counts from all libraries in each condition, and using a dense ranking.

#### Integration over all data types and experiments

Different experiments and techniques might have different power and resolution at ranking genes in a same condition. For instance, some *in situ* hybridization data might allow to rank 5 genes in a condition; while RNA-Seq data in the same condition might allow to rank 30,000 genes. To correct for this, we normalize the mean or dense ranks for each gene in each condition and for each data type, based on the max rank for this data type in this condition, as compared to the max rank over all conditions and data types, independently for each species (see eq. 4). With the information normalized per data type we compute a global weighted mean rank for each gene and condition. All data types are used for ranks shown on the Gene Page, although it is possible to use only some if needed. This mean is computed by using the normalized mean ranks or normalized dense ranks for a gene in a condition, from each data type considered, and as weights, the sum of distinct rank counts for RNA-Seq and Affymetrix data, the max rank in each condition for EST and *in situ* hybridization data (see eq. 5).

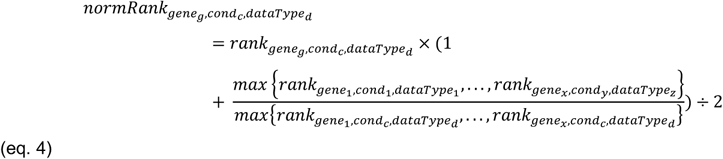

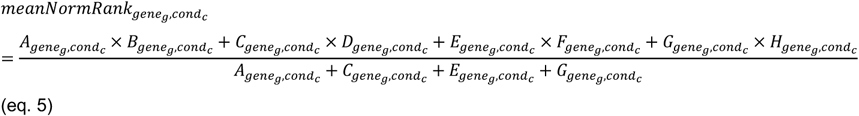

where 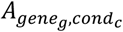 is the sum of the counts of distinct ranks in the RNA-Seq libraries in *cond*_*c*_ studying *gene*_*g*_, 0 if there is no RNA-Seq data for this gene in this condition; 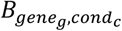 is the mean normalized rank of *gene*_*g*_ over the RNA-Seq libraries in *cond*_*c*_ with data for *gene*_*g*_; 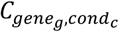 is the sum of the counts of distinct ranks in the Affymetrix chips in *cond*_*c*_ studying *gene*_*g*_, 0 if there is no Affymetrix data for this gene in this condition; 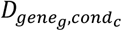 is the mean normalized rank of *gene*_*g*_ over the Affymetrix chips in *cond*_*c*_ with data for *gene*_*g*_; 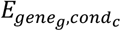 is the max dense rank from *in situ* hybridization data in *cond*_*c*_ over all genes, 0 if there is no *in situ* hybridization data for *gene*_*g*_ in this condition; 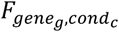 is the normalized dense rank of *gene*_*g*_ from *in situ* hybridization data in *cond*_*c*_; 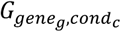 is the max dense rank from EST data in *cond*_*c*_ over all genes, 0 if there is no EST data for *cond*_*c*_ in this condition; 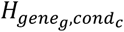 is the normalized dense rank of *gene*_*g*_ from EST data in *cond*_*c*_;

We transform global weighted mean ranks into expression scores. To compute the expression score of a gene in a condition, we retrieve the max rank over all conditions and data types considered, for this species; and the global weighted mean rank for a gene in a condition, computed over all data types considered and data available, as described above. The expression score is computed according to eq. 6.

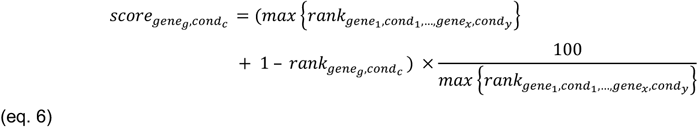

## RESULTS: USING THE BGEE RESOURCE

We describe here: i) the data available in Bgee that we have produced; ii) the web interfaces to leverage these data for biological insights; iii) the resources to access Bgee data (annotated and processed expression values, calls of expression).

### Overview of Bgee

One of the advantages of the integrative approach of Bgee is to obtain data for as many conditions as possible, with as much anatomical and developmental detail as possible. Moreover, new technologies do not require a new database, but allow integration with historical data.

For developing Bgee, we have curated and reprocessed all expression data to the same standard, of high quality, non-redundant, healthy wild-type data. This is a first level of data integration.

Bgee provides a single answer to the question “where and when is this gene expressed?”. For instance, for the insulin gene, both human and mouse insulin have in Bgee top expression in islets of Langerhans and more generally pancreas; in human the information is even more fine-grained, with top expression in the Beta cells. Users of this second level of integration (calls) are free to ignore the complexity of the underlying data, to concentrate on the biological signal of interest, while we also provide high quality processed data for downstream studies.

### Data provided by Bgee

In this section we describe the Bgee release 14.1 datasets of: i) annotated and processed expression values; ii) calls of presence/absence of expression and expression ranks; iii) anatomical and developmental stage similarity information. All original data sources used to build Bgee release 14.1 are in Table 1. All data are published under the Creative Commons Zero license (CC0).

#### Annotated and processed expression values

Bgee 14 includes 29 animal species. Anatomical localization is annotated to the multi-species anatomical ontology Uberon (17), and developmental stages to a multi-species ontology (Table S1), which integrates existing ontologies when available. Sex and strain are annotated with a basic controlled vocabulary, and species with the NCBI taxonomy(13). We have curated all data to retain only healthy, wild-type data: no treatment, no gene knockout, etc. Since we have annotated and remapped all these data in a consistent framework using ontologies, and reprocessed all of them, it allows a complete integration of all data. Bgee thus allows users to retrieve a dataset of expression data consistently curated, annotated, and processed, usable in their own downstream analyses.

Statistics about the data integrated per species are presented in Table 2.

**Table 2:** Data statistics for release Bgee 14.1 per species for the 29 species included in this version. This table provides the number of EST libraries integrated, number of *in situ* hybridization experiments and evidence (e.g., the image of a staining) counts, number of Affymetrix chips and experiments, number of RNA-Seq libraries and experiments.

| Species | EST | In Situ |  | Affymetrix |  | RNA-Seq |  |
| --- | --- | --- | --- | --- | --- | --- | --- |
|  | libraries | evidence | experiments | chips | experiments | libraries | experiments |
| Anolis carolinensis | - | - | - | - | - | 31 | 5 |
| Bos taurus | - | - | - | - | - | 121 | 8 |
| Caenorhabditi selegans | - | 360 | 141 | 177 | 34 | 41 | 6 |
| Canis lupus familiaris | - | - | - | - | - | 141 | 21 |
| Cavia porcellus | - | - | - | - | - | 28 | 4 |
| Danio rerio | 108 | 42153 | 4674 | 186 | 33 | 147 | 16 |
| Drosophila ananassae | - | - | - | - | - | 4 | 1 |
| Drosophila melanogaster | 62 | 92227 | 5468 | 965 | 92 | 253 | 11 |
| Drosophila mojavensis | - | - | - | - | - | 8 | 1 |
| Drosophila pseudoobscura | - | - | - | - | - | 10 | 1 |
| Drosophila simulans | - | - | - | - | - | 15 | 2 |
| Drosophila virilis | - | - | - | - | - | 4 | 1 |
| Drosophila yakuba | - | - | - | - | - | 4 | 1 |
| Equus caballus | - | - | - | - | - | 232 | 22 |
| Erinaceus europaeus | - | - | - | - | - | 6 | 1 |
| Felis catus | - | - | - | - | - | 32 | 5 |
| Gallus gallus | - | - | - | - | - | 48 | 6 |
| Gorilla gorilla | - | - | - | - | - | 13 | 2 |
| Homo sapiens | 2393 | - | - | 5452 | 323 | 5676 | 36 |
| Macaca mulatta | - | - | - | 14 | 1 | 238 | 18 |
| Monodelphis domestica | - | - | - | - | - | 108 | 15 |
| Mus musculus | 706 | 223513 | 37654 | 6095 | 698 | 330 | 30 |
| Ornithorhynchus anatinus | - | - | - | - | - | 21 | 4 |
| Oryctolagus cuniculus | - | - | - | - | - | 55 | 13 |
| Pan paniscus | - | - | - | - | - | 13 | 2 |
| Pan troglodytes | - | - | - | - | - | 250 | 18 |
| Rattus norvegicus | - | - | - | 107 | 2 | 106 | 8 |
| Sus scrofa | - | - | - | - | - | 169 | 14 |
| Xenopus tropicalis | 66 | 2400 | 1304 | - | - | 259 | 5 |

##### RNA-Seq data

For RNA-Seq data, we have remapped all raw data to transcriptome to produce counts at the transcript level, then aggregated them per gene. These aggregated counts are used to compute FPKMs and TPMs per gene. We have notably re-curated the GTEx human dataset phs000424.v6.p1(7). Only 50% of samples were kept, to discard unhealthy or contaminated samples (see Table S1 for link to GTEx criteria), representing a high quality subset of GTEx. For each RNA-Seq library, we provide detailed information, including annotations, library information, and relevant statistics (see online documentation). For each gene in each library, we provide several measures of expression level, and calls of presence/absence of expression (available in download TSV files).

##### Affymetrix data

For Affymetrix data, we have processed raw CEL files when available, or used MAS5 processed files when raw CEL files were not available. Similar to RNA-Seq libraries, for each Affymetrix chip, we provide annotations, chip information, and relevant statistics. In Affymetrix, expression is measured per probeset rather than per gene; several probesets can map to one gene, and provide different measures. For each probeset in each chip, we provide the gene the probeset maps to, the signal intensity, and the call of presence/absence of expression (see online documentation) (available in download TSV files).

##### *In situ* hybridization data

We have retrieved *in situ* hybridization data from relevant Model Organism Databases (MODs; Table 1). For each evidence, we do not store the original image, or paper figure, but we provide a link to the original data. For each evidence and each spot, meaning the report of an area with staining from expression of a gene, or lack of staining from absence of expression of a gene, we provide annotations, mapping to gene, call of presence/absence of expression and quality of the call for this spot. At present, these data are not available for direct download, but are used in integrated calls and scores.

##### Expressed Sequence Tags

Both EST databases which we used as data sources (Table 1) are now retired, thus we are no longer updating these data. For each EST library, we provide annotations. We provide the mapping between genes and ESTs per library. For each gene in each library, we provide the number of ESTs mapped to it, and the call of presence of expression. Despite the growth in other data, there are still conditions where ESTs provide the only direct observation. For example, mouse insulin-1 has only EST data for pancreas at 1 week. Note that calls of absence of expression are not produced from EST data. At present, these data are not available for direct download, but are used in integrated calls and scores.

#### Calls of presence/absence of expression and expression ranks

##### Expression calls

A call corresponds to a unique combination of a gene in a condition, with reported presence or absence of expression. In Bgee 14, only conditions combining species, anatomical entity, and life stage are available to users of our calls; information about sex and strain will be publicly available for calls in a future release. We produce calls for each evidence (e.g., RNA-Seq library, Affymetrix chip) integrated into Bgee. For continuous data types, we apply a threshold which depends on the data type: for Affymetrix a Wilcoxon test on the signal of the probesets against a subset of weakly expressed probesets(49); for RNA-seq, a library-specific threshold depending on the distribution of TPMs on intergenic sequences (Julien Roux, Marta Rosikiewicz, Julien Wollbrett, Sara S. Fonseca Costa, Marc Robinson-Rechavi, Frederic B. Bastian; in preparation); and for ESTs a threshold based on the number of tags(50). These calls can then be integrated over experiments and over data types.

We have propagated these individual calls along a graph of conditions (see *Materials and Methods*). Then, we have integrated these individual calls from multiple experiments, propagated along the condition graph, to produce one global call of presence/absence of expression per gene - condition, associated to a confidence level(39). As a result, for each gene in each condition, Bgee provides one global call, the confidence level of the call (gold, silver, bronze), the number of experiments from each data type supporting the presence and absence of expression, and whether the call has been propagated or observed directly in the condition (available in TSV download files).

##### Expression ranks and scores

To compare expression between experiments in a quantitative manner, we rank genes in a condition based on their expression level. We have also integrated these ranks over all data types and experiments. The lower the rank score of a gene in a condition, the higher its expression. Because this is not very intuitive to users, we also compute an “expression score” for visualisation: the top expressed gene has a rank score of 1 and an expression score of 100, and the lowest expressed has the maximum rank and an expression score of 100/Max(ranks). Expression ranks can be retrieved per anatomical entity and developmental stage.

#### Anatomical and developmental similarity

We use the ontology of homology and related concepts(51) to capture the type of similarity between anatomical entities. These similarity annotations can be retrieved from GitHub (full link in Table S1), and will be described in detail elsewhere (Anne Niknejad, Marc Robinson-Rechavi, Frederic B. Bastian; unpublished). It should be emphasized that they are derived from primary literature (e.g., paleontology, Evo-Devo), not from the expression data in Bgee; thus the anatomical homology annotations and the gene expression calls are independent. As of Bgee 14.1, we have integrated 2,328 relations of homology, involving 1,845 anatomical entities.

For development stages, we have merged all the developmental stage ontologies of species integrated in Bgee into one common multi-species ontology, within Uberon. Common general stages have been merged between species, and more precise species-specific stages are children of these general terms. For instance, the human-specific precise stage HsapDv:0000016 “Carnegie stage 09” is a child of the more general, multi-species term UBERON:0000111 “organogenesis stage”. While we could not map data to the exact equivalent of Carnegie stage 09 in all species, we could compare them at the more general organogenesis stage, thanks to propagation of calls to parent terms. This makes comparison with a developmental aspect possible.

### Web interface of Bgee

The Bgee Gene page (Figure 3) provides for each gene the conditions where it is expressed, sorted by their expression rank. Primary sorting is on anatomical entities.

**Figure 3:**
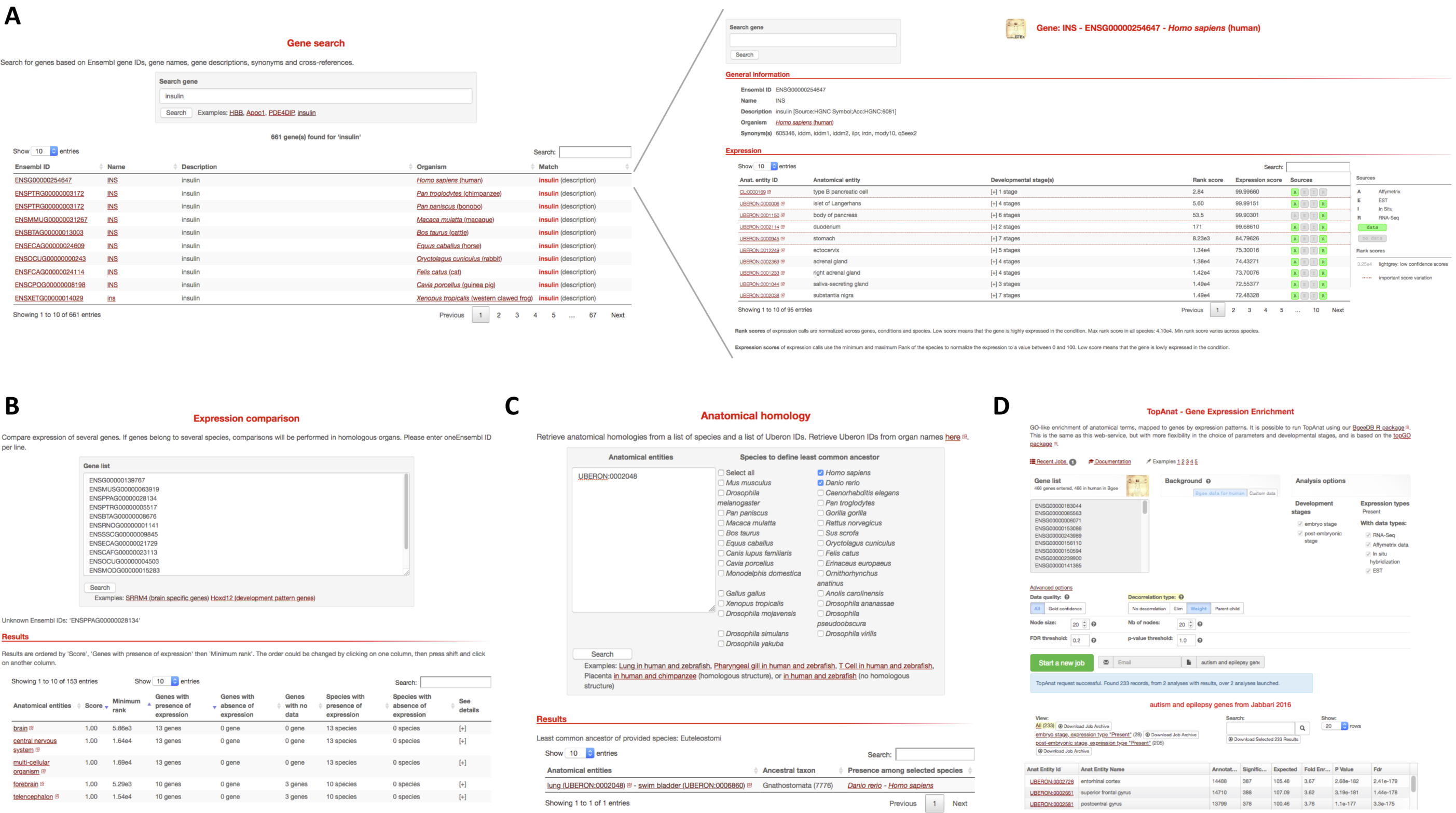
screenshots of the Bgee web interfaces. A: example of gene search (top left) for the term “insulin” (https://bgee.org/?page=gene&query=insulin), allowing to go to the gene page (top right) displaying ranked conditions with expression for the human gene INS (https://bgee.org/?page=gene&gene_id=ENSG00000254647). B: example of comparison of expression patterns for the SRRM4 genes (brain-related genes) in 13 species (https://bgee.org/?page=expression_comparison&data=34beddfc93bb7fbb440e757e6de24d91fc0ce177). C: Anatomical homology retrieval tool, with here an example query allowing to identify swim bladder as the anatomical structure in zebrafish homlogous to the human lung (https://bgee.org/?page=anat_similarities&species_list=9606&species_list=7955&ae_list=UBERON%3A0002048). D: example of TopAnat analysis on a set of human genes associated to autism and epilepsy, identifying the enriched conditions with expression of these genes as specific brain regions (https://bgee.org/?page=top_anat#/result/8fce889da7b4519c5792573ed3933032c8122819/).

The anatomical expression enrichment test TopAnat (Figure 3) uses a similar approach to Gene Ontology enrichment tests(52)(53), but genes are associated to the anatomical structures from Uberon by their expression calls, instead of to their functional classification (Roux J., Seppey M., Sanjeev K., Rech de Laval V., Moret P., Artimo P., Duvaud S., Ioannidis V., Stockinger H., Robinson-Rechavi M., Bastian F.B.; unpublished). The algorithms from the package topGO(54) are available in TopAnat to account for the non-independence of anatomical structures, and avoid the over-representation of less-informative top-level terms. As an example, we have used TopAnat to analyze a list of genes associated with autism and epilepsy in human(55). TopAnat returns a list of anatomical structures where expression of these genes is enriched relative to the background of all human genes with expression. These structures are almost all specific brain regions, including all brain regions known to be affected by autism(56): frontotemporal lobe (examples of TopAnat results part of this structure: UBERON:0002771 “middle temporal gyrus” and UBERON:0002810 “right frontal lobe”); frontoparietal cortex (UBERON:0001872 “parietal lobe” and UBERON:0001870 “frontal cortex”); amygdala (UBERON:0001876 “amygdala”); hippocampus (UBERON:0003881 “CA1 field of hippocampus”); basal ganglia (UBERON:0002038 “substantia nigra”, part of UBERON:0002420 “basal ganglion”); and anterior cingulate cortex (UBERON:0009835 “anterior cingulate cortex”). Of note, TopAnat can be slow because of propagation of calls in the anatomical ontology, which is redone on the fly by the topGO package to allow decorrelation. TopAnat is available as a web-tool, and in the BgeeDB R package (see below).

The Bgee website makes it possible to compare expression patterns between genes, within and between species. When genes in a single species are compared, Expression Comparison (see figure 3) considers present/absent expression calls in anatomical entities (e.g., “lung” in human). When genes from several species are compared, it considers calls in homologous anatomical entities (e.g., “lung - swim bladder” for comparing data in human and zebrafish(57–59)). For each of these entities, it then displays the genes from the user list that are either expressed, or not expressed, or have no data. It offers different sorting methods of these entities, by expression rank and by conservation of calls.

For instance, when studying the conservation of expression of the brain specific genes SRRM4 in Tetrapoda, the top homologous anatomical entities, as of Bgee 14.1, when sorted by maximum expression conservation and minimum expression rank, are “forebrain”, “telencephalon”, and “cerebral hemisphere” (https://bgee.org/bgee14_1/?page=expression_comparison&data=34beddfc93bb7fbb440e757e6de24d91fc0ce177). Relations of historical homology can also be queried directly through the homology retrieval tool (see figure 3). Thus they can be used for other applications than expression in Bgee, such as phenotypes. For instance, for comparing such anatomy-related data, a user may ask: “what is, in human, the comparable organ to the zebrafish ‘pharyngeal gill’?”. The homology retrieval tool will return the anatomical entity in human “parathyroid gland” (https://bgee.org/bgee14_1/?page=anat_similarities&species_list=9606&species_list=7955&ae_list=UBERON%3A0000206). Indeed, the parathyroid gland and the pharyngeal gill likely derive from a common ancestral structure, present in the ancestor of Vertebrata, as they both regulate extracellular calcium levels, and the parathyroid gland is positioned within the pharynx in Tetrapoda(60). We have captured this information in our annotations of similarities between anatomical entities, that the homology retrieval webtool uses to answer the user query.

### Resources to access data

We provide access to the annotated and processed expression values through the Bioconductor(61) R(62) package BgeeDB(63). BgeeDB is a collection of functions to import into R these data, facilitating downstream analyses. BgeeDB also allows to run TopAnat analyses, offering more flexibility in the choice of input data and analysis parameters than the web-interface.

Bgee also provides a SPARQL(64) endpoint which is based on the EasyBgee database. EasyBgee is a lighter version of the Bgee database, that contains the most useful information, made explicit. The endpoint is accessible at the address https://bgee.org/sparql/. In addition to SPARQL endpoint querying through any programmatic language, a web-interface also offers the possibility to run more user-friendly queries, developed as part of the BioSODA project(65). This web-interface is available at http://biosoda.expasy.org/. Bgee specific queries are present under the category “Bgee database queries”.

We provide TSV files to retrieve, for each species: i) annotated and processed expression values for RNA-Seq and Affymetrix data; and ii) calls of presence/absence of expression with confidence and rank scores generated from all data types.

## DISCUSSION

The philosophy of Bgee is to provide information and data which users can trust, so that they can be built on to do further work. Expert manual curation is at the core of providing trustworthy information (66–69). It is key to certify that expression data is from healthy wild-type samples, and to annotate precisely and accurately. It is also the only way to generate reliable annotations of anatomical homology. While expression data analysis is not specific to Bgee(8), our approach is to focus on delivering clear biological signals, and choosing the best methodology to do so. For users to have an easy and intuitive access to information from expression data, despite its complexity, we have chosen to present different views which answer different questions. These include summarized views, such as the Gene Page or Expression Comparison, with more advanced access, such as the R packages or SPARQL endpoint, to serve different use cases and ensure flexibility.

Comparative transcriptomics is essential to understand the molecular basis for phenotypes, such as, for instance, evolution of animal morphology(70), embryonic cell differentiation(71), species lifespan(72, 73), or cancer evolution(74). Bgee, by generating comparable reference sets of expression patterns in multiple species, and homology relations to link them, is the first resource to allow the systematic and automated comparison of gene expression patterns between species.

These comparisons are based on the present/absent calls of expression produced by Bgee. It is thus essential that these calls capture the relevant functional aspects of gene expression. This relevance is demonstrated, for instance, by the results provided by our webtools “Gene Page”, and by TopAnat.

On the Gene Page, the top ranked conditions of genes are relevant to their known biology (e.g., as of Bgee 14.1: several muscle regions for human *PDE4DIP* gene, “liver” for mouse *Apoc1* gene, “pancreas” for Xenopus *ins* gene). In TopAnat, results for list of genes are highly representative of their known function (e.g., as of Bgee 14.1: top ranked condition is “spermatocyte” for a list of mouse genes associated with spermatogenesis, “musculature of body” for a list of cow genes with a relation to muscle in their description). Bgee thus provides reference sets of expression patterns that are accurate and predictive of gene functions.

Here we have presented the latest release of Bgee, which integrates expression information from four well established data types. An important feature is that the model of Bgee allows integration of new data types into the same framework. From each new data type, we need to define quality control criteria, conditions for calling gene expression present or absent, and rules for expression ranks. Once this is done, our model will allow views and tools such as the Gene Page, the Expression Comparison, or TopAnat to make use of the new data together with the previously available data. Notably, single-cell RNA-Seq presents an important perspective of combining anatomical precision beyond that of in situ hybridization with the broad coverage of microarrays or RNA-Seq. Unlike dedicated single-cell databases, Bgee offers the perspective of a unified view of gene expression from the cellular to the organism level, which we believe will be increasingly relevant.

## AVAILABILITY

Bgee, documentation, and links to resources used, are available at https://bgee.org/. Source code and annotations are available in the GitHub repository (https://github.com/BgeeDB). R packages are available in Bioconductor (https://bioconductor.org/packages/BgeeDB/ and https://bioconductor.org/packages/BgeeCall/).

## SUPPLEMENTARY DATA

Supplementary Data are available at NAR online.

## ACKNOWLEDGMENT

Chris Mungall, Melissa Haendel, Suzanna Lewis, David Osumi-Sutherland, Daniela Raciti, and Torsten Henrich helped and coordinated with us concerning ontologies and annotation. Vincent Laudet was instrumental in helping the project get started. Bgee was also improved by very helpful discussions with Amos Bairoch, Pascale Gaudet, Lydie Lane, Christophe Dessimoz, Adrian Altenhoff, Natasha Glover, and many members of the SIB Swiss Institute of Bioinformatics and SIB UX team; as well as many members of the biocuration and OBO Foundry communities, notably the teams of the Uberon project, the Model Organisms Databases BDGP, FlyBase, GXD, WormBase, Xenbase, and ZFIN, as well as the Alliance for Genome Resources (AGR), the ArrayExpress team, and the owlcollab and obophenotype communities. The NCBI Taxonomy rapidly added new taxa when needed for homology annotations.

The Genotype-Tissue Expression (GTEx) Project was supported by the <u>Common Fund</u> of the Office of the Director of the National Institutes of Health, and by NCI, NHGRI, NHLBI, NIDA, NIMH, and NINDS. The data integrated in Bgee described in this manuscript were obtained from dbGaP accession number phs000424.v6.p1.

Finally, we thank all members and alumni of the Robinson-Rechavi lab; special thanks to Nadezda Kryuchkova for discussions on tissue-specificity, and to Andrea Komljenovic and Tina Begum for suggestions of data to annotate and for use-cases.

## REFERENCES

1. Haeussler, M., Zweig, A.S., Tyner, C., Speir, M.L., Rosenbloom, K.R., Raney, B.J., Lee, C.M., Lee, B.T., Hinrichs, A.S., Gonzalez, J.N., et al. (2018) The UCSC Genome Browser database: 2019 update. Nucleic Acids Res., 47, D853–D858.

2. Howe, K.L., Contreras-Moreira, B., De Silva, N., Maslen, G., Akanni, W., Allen, J., Alvarez-Jarreta, J., Barba, M., Bolser, D.M., Cambell, L., et al. (2019) Ensembl Genomes 2020-enabling non-vertebrate genomic research. Nucleic Acids Res., 10.1093/nar/gkz890.

3. Yates, A.D., Achuthan, P., Akanni, W., Allen, J., Allen, J., Alvarez-Jarreta, J., Amode, M.R., Armean, I.M., Azov, A.G., Bennett, R., et al. (2019) Ensembl 2020. Nucleic Acids Res., 10.1093/nar/gkz966.

4. Roux, J., Rosikiewicz, M. and Robinson-Rechavi, M. (2015) What to compare and how: Comparative transcriptomics for Evo-Devo: COMPARATIVE TRANSCRIPTOMICS FOR Evo-Devo. J. Exp. Zoolog. B Mol. Dev. Evol., 324, 372–382.

5. Brown, G.R., Hem, V., Katz, K.S., Ovetsky, M., Wallin, C., Ermolaeva, O., Tolstoy, I., Tatusova, T., Pruitt, K.D., Maglott, D.R., et al. (2014) Gene: a gene-centered information resource at NCBI. Nucleic Acids Res., 43, D36–D42.

6. Fagerberg, L., Hallström, B.M., Oksvold, P., Kampf, C., Djureinovic, D., Odeberg, J., Habuka, M., Tahmasebpoor, S., Danielsson, A., Edlund, K., et al. (2014) Analysis of the human tissue-specific expression by genome-wide integration of transcriptomics and antibody-based proteomics. Mol. Cell. Proteomics MCP, 13, 397–406.

7. Carithers, L.J., Ardlie, K., Barcus, M., Branton, P.A., Britton, A., Buia, S.A., Compton, C.C., DeLuca, D.S., Peter-Demchok, J., Gelfand, E.T., et al. (2015) A Novel Approach to High-Quality Postmortem Tissue Procurement: The GTEx Project. Biopreservation Biobanking, 13, 311–319.

8. Rung, J. and Brazma, A. (2012) Reuse of public genome-wide gene expression data. Nat. Rev. Genet., 14, 89.

9. Papatheodorou, I., Fonseca, N.A., Keays, M., Tang, Y.A., Barrera, E., Bazant, W., Burke, M., Füllgrabe, A., Fuentes, A.M.-P., George, N., et al. (2017) Expression Atlas: gene and protein expression across multiple studies and organisms. Nucleic Acids Res., 46, D246–D251.

10. Palasca, O., Santos, A., Stolte, C., Gorodkin, J. and Jensen, L.J. (2018) TISSUES 2.0: an integrative web resource on mammalian tissue expression. Database, 2018.

11. Vize, P.D. and Westerfield, M. (2015) Model organism databases. genesis, 53, 449–449.

12. Howe, D.G., Blake, J.A., Bradford, Y.M., Bult, C.J., Calvi, B.R., Engel, S.R., Kadin, J.A., Kaufman, T.C., Kishore, R., Laulederkind, S.J.F., et al. (2018) Model organism data evolving in support of translational medicine. Lab Anim., 47, 277–289.

13. Federhen, S. (2011) The NCBI Taxonomy database. Nucleic Acids Res., 40, D136–D143.

14. Ashburner, M., Ball, C.A., Blake, J.A., Botstein, D., Butler, H., Cherry, J.M., Davis, A.P., Dolinski, K., Dwight, S.S., Eppig, J.T., et al. (2000) Gene ontology: tool for the unification of biology. The Gene Ontology Consortium. Nat. Genet., 25, 25–29.

15. The Gene Ontology Consortium (2019) The Gene Ontology Resource: 20 years and still GOing strong. Nucleic Acids Res., 47, D330–D338.

16. Altenhoff, A.M., Glover, N.M., Train, C.-M., Kaleb, K., Warwick Vesztrocy, A., Dylus, D., de Farias, T.M., Zile, K., Stevenson, C., Long, J., et al. (2018) The OMA orthology database in 2018: retrieving evolutionary relationships among all domains of life through richer web and programmatic interfaces. Nucleic Acids Res., 46, D477–D485.

17. Haendel, M.A., Balhoff, J.P., Bastian, F.B., Blackburn, D.C., Blake, J.A., Bradford, Y., Comte, A., Dahdul, W.M., Dececchi, T.A., Druzinsky, R.E., et al. (2014) Unification of multi-species vertebrate anatomy ontologies for comparative biology in Uberon. J. Biomed. Semant., 5, 21.

18. Hayamizu, T.F., Baldock, R.A. and Ringwald, M. (2015) Mouse anatomy ontologies: enhancements and tools for exploring and integrating biomedical data. Mamm. Genome, 26, 422–430.

19. Costa, M., Reeve, S., Grumbling, G. and Osumi-Sutherland, D. (2013) The Drosophila anatomy ontology. J. Biomed. Semant., 4, 32.

20. Van Slyke, C.E., Bradford, Y.M., Westerfield, M. and Haendel, M.A. (2014) The zebrafish anatomy and stage ontologies: representing the anatomy and development of Danio rerio. J. Biomed. Semant., 5, 12.

21. Segerdell, E., Bowes, J.B., Pollet, N. and Vize, P.D. (2008) An ontology for Xenopus anatomy and development. BMC Dev. Biol., 8, 92.

22. Segerdell, E., Ponferrada, V.G., James-Zorn, C., Burns, K.A., Fortriede, J.D., Dahdul, W.M., Vize, P.D. and Zorn, A.M. (2013) Enhanced XAO: the ontology of Xenopus anatomy and development underpins more accurate annotation of gene expression and queries on Xenbase. J. Biomed. Semant., 4, 31.

23. Harris, T.W., Arnaboldi, V., Cain, S., Chan, J., Chen, W.J., Cho, J., Davis, P., Gao, S., Grove, C.A., Kishore, R., et al. (2019) WormBase: a modern Model Organism Information Resource. Nucleic Acids Res., 10.1093/nar/gkz920.

24. Wang, J., Zou, H., Chen, L., Long, X., Lan, J., Liu, W., Ma, L., Wang, C., Xu, X., Ren, L., et al. (2017) Convergent and divergent genetic changes in the genome of Chinese and European pigs. Sci. Rep., 7, 8662.

25. Kozomara, A., Birgaoanu, M. and Griffiths-Jones, S. (2019) miRBase: from microRNA sequences to function. Nucleic Acids Res., 47, D155–D162.

26. Barrett, T., Wilhite, S.E., Ledoux, P., Evangelista, C., Kim, I.F., Tomashevsky, M., Marshall, K.A., Phillippy, K.H., Sherman, P.M., Holko, M., et al. (2012) NCBI GEO: archive for functional genomics data sets—update. Nucleic Acids Res., 41, D991–D995.

27. Tryka, K.A., Hao, L., Sturcke, A., Jin, Y., Wang, Z.Y., Ziyabari, L., Lee, M., Popova, N., Sharopova, N., Kimura, M., et al. (2013) NCBI’s Database of Genotypes and Phenotypes: dbGaP. Nucleic Acids Res., 42, D975–D979.

28. Kodama, Y., on behalf of the International Nucleotide Sequence Database Collaboration, Shumway, M., on behalf of the International Nucleotide Sequence Database Collaboration, Leinonen, R. and on behalf of the International Nucleotide Sequence Database Collaboration (2011) The sequence read archive: explosive growth of sequencing data. Nucleic Acids Res., 40, D54–D56.

29. Athar, A., Füllgrabe, A., George, N., Iqbal, H., Huerta, L., Ali, A., Snow, C., Fonseca, N.A., Petryszak, R., Papatheodorou, I., et al. (2019) ArrayExpress update - from bulk to single-cell expression data. Nucleic Acids Res., 47, D711–D715.

30. Hammonds, A.S., Bristow, C.A., Fisher, W.W., Weiszmann, R., Wu, S., Hartenstein, V., Kellis, M., Yu, B., Frise, E. and Celniker, S.E. (2013) Spatial expression of transcription factors in Drosophila embryonic organ development. Genome Biol., 14, R140.

31. Tomancak, P., Berman, B.P., Beaton, A., Weiszmann, R., Kwan, E., Hartenstein, V., Celniker, S.E. and Rubin, G.M. (2007) Global analysis of patterns of gene expression during Drosophila embryogenesis. Genome Biol., 8, R145.

32. Tomancak, P., Beaton, A., Weiszmann, R., Kwan, E., Shu, S., Lewis, S.E., Richards, S., Ashburner, M., Hartenstein, V., Celniker, S.E., et al. (2002) Systematic Determination of Patterns of Gene Expression During Drosophila Embryogenesis. Genome Biol., 3, research0088.1.

33. Thurmond, J., Goodman, J.L., Strelets, V.B., Attrill, H., Gramates, L.S., Marygold, S.J., Matthews, B.B., Millburn, G., Antonazzo, G., Trovisco, V., et al. (2018) FlyBase 2.0: the next generation. Nucleic Acids Res., 47, D759–D765.

34. Smith, C.M., Hayamizu, T.F., Finger, J.H., Bello, S.M., McCright, I.J., Xu, J., Baldarelli, R.M., Beal, J.S., Campbell, J., Corbani, L.E., et al. (2018) The mouse Gene Expression Database (GXD): 2019 update. Nucleic Acids Res., 47, D774–D779.

35. Karimi, K., Fortriede, J.D., Lotay, V.S., Burns, K.A., Wang, D.Z., Fisher, M.E., Pells, T.J., James-Zorn, C., Wang, Y., Ponferrada, V.G., et al. (2017) Xenbase: a genomic, epigenomic and transcriptomic model organism database. Nucleic Acids Res., 46, D861–D868.

36. Howe, D.G., Bradford, Y.M., Conlin, T., Eagle, A.E., Fashena, D., Frazer, K., Knight, J., Mani, P., Martin, R., Moxon, S.A.T., et al. (2012) ZFIN, the Zebrafish Model Organism Database: increased support for mutants and transgenics. Nucleic Acids Res., 41, D854–D860.

37. Landgraf, P., Rusu, M., Sheridan, R., Sewer, A., Iovino, N., Aravin, A., Pfeffer, S., Rice, A., Kamphorst, A.O., Landthaler, M., et al. (2007) A Mammalian microRNA Expression Atlas Based on Small RNA Library Sequencing. Cell, 129, 1401–1414.

38. Pontius, J.U., Wagner, L. and Schuler, G.D. (2004) UniGene: A unified view of the transcriptome. In: The NCBI Handbook National Center for Biotechnology Information.

39. Bastian, F.B., Chibucos, M.C., Gaudet, P., Giglio, M., Holliday, G.L., Huang, H., Lewis, S.E., Niknejad, A., Orchard, S., Poux, S., et al. (2015) The Confidence Information Ontology: a step towards a standard for asserting confidence in annotations. Database, 2015.

40. Rosikiewicz, M., Comte, A., Niknejad, A., Robinson-Rechavi, M. and Bastian, F.B. (2013) Uncovering hidden duplicated content in public transcriptomics data. Database, 2013.

41. Bray, N.L., Pimentel, H., Melsted, P. and Pachter, L. (2016) Near-optimal probabilistic RNA-seq quantification. Nat. Biotechnol., 34, 525.

42. Rosikiewicz, M. and Robinson-Rechavi, M. (2016) IQRray, a new method for Affymetrix microarray quality control, and the homologous organ conservation score, a new benchmark method for quality control metrics. Bioinformatics, 32, 2565–2565.

43. Wu, Z., Irizarry, R.A., Gentleman, R., Martinez-Murillo, F. and Spencer, F. (2004) A Model-Based Background Adjustment for Oligonucleotide Expression Arrays. J. Am. Stat. Assoc., 99, 909–917.

44. Hubbell, E., Liu, W.-M. and Mei, R. (2002) Robust estimators for expression analysis. Bioinformatics, 18, 1585–1592.

45. Altschul, S.F., Gish, W., Miller, W., Myers, E.W. and Lipman, D.J. (1990) Basic local alignment search tool. J. Mol. Biol., 215, 403–410.

46. Kalderimis, A., Lyne, R., Butano, D., Contrino, S., Lyne, M., Heimbach, J., Hu, F., Smith, R., Štěpán, R., Sullivan, J., et al. (2014) InterMine: extensive web services for modern biology. Nucleic Acids Res., 42, W468–W472.

47. Liu, W. -m, Mei, R., Di, X., Ryder, T.B., Hubbell, E., Dee, S., Webster, T.A., Harrington, C.A., Ho, M. -h., Baid, J., et al. (2002) Analysis of high density expression microarrays with signed-rank call algorithms. Bioinformatics, 18, 1593–1599.

48. Choe, S.E., Boutros, M., Michelson, A.M., Church, G.M. and Halfon, M.S. (2005) Preferred analysis methods for Affymetrix GeneChips revealed by a wholly defined control dataset. Genome Biol., 6, R16.

49. Schuster, E.F., Blanc, E., Partridge, L. and Thornton, J.M. (2007) Correcting for sequence biases in present/absent calls. Genome Biol., 8, R125.

50. Audic, S. and Claverie, J.-M. (1997) The Significance of Digital Gene Expression Profiles. Genome Res., 7, 986–995.

51. Roux, J. and Robinson-Rechavi, M. (2010) An ontology to clarify homology-related concepts. Trends Genet., 26, 99–102.

52. Yon Rhee, S., Wood, V., Dolinski, K. and Draghici, S. (2008) Use and misuse of the gene ontology annotations. Nat. Rev. Genet., 9, 509–515.

53. Dessimoz, C. and Škunca, N. (2017) The Gene Ontology Handbook Humana Press New York, NY, USA:

54. Alexa, A., Rahnenfuhrer, J. and Lengauer, T. (2006) Improved scoring of functional groups from gene expression data by decorrelating GO graph structure. Bioinformatics, 22, 1600–1607.

55. Jabbari, K. and Nürnberg, P. (2016) A genomic view on epilepsy and autism candidate genes. Genomics, 108, 31–36.

56. Ha, S., Sohn, I.-J., Kim, N., Sim, H.J. and Cheon, K.-A. (2015) Characteristics of Brains in Autism Spectrum Disorder: Structure, Function and Connectivity across the Lifespan. Exp. Neurobiol., 24, 273–284.

57. Schmidt-Rhaesa, A. (2007) The evolution of organ systems Oxford University Press, Oxford?; New York.

58. Zheng, W., Wang, Z., Collins, J.E., Andrews, R.M., Stemple, D. and Gong, Z. (2011) Comparative Transcriptome Analyses Indicate Molecular Homology of Zebrafish Swimbladder and Mammalian Lung. PLoS ONE, 6, e24019.

59. Zaccone, D., Sengar, M., Lauriano, E.R., Pergolizzi, S., Macri’, F., Salpietro, L., Favaloro, A., Satora, L., Dabrowski, K. and Zaccone, G. (2012) Morphology and innervation of the teleost physostome swim bladders and their functional evolution in non-teleostean lineages. Acta Histochem., 114, 763–772.

60. Graham, A., Okabe, M. and Quinlan, R. (2005) The role of the endoderm in the development and evolution of the pharyngeal arches: Endoderm in the development and evolution of the pharyngeal arches, A. Graham et al. J. Anat., 207, 479–487.

61. Huber, W., Carey, V.J., Gentleman, R., Anders, S., Carlson, M., Carvalho, B.S., Bravo, H.C., Davis, S., Gatto, L., Girke, T., et al. (2015) Orchestrating high-throughput genomic analysis with Bioconductor. Nat. Methods, 12, 115–121.

62. R Core Team (2018) R: A Language and Environment for Statistical Computing R Foundation for Statistical Computing, Vienna, Austria.

63. Komljenovic, A., Roux, J., Wollbrett, J., Robinson-Rechavi, M. and Bastian, F.B. (2018) BgeeDB, an R package for retrieval of curated expression datasets and for gene list expression localization enrichment tests. F1000Research, 5, 2748.

64. Segaran, T., Taylor, J. and Evans, C. (2009) Programming the Semantic Web 1st ed. O’Reilly, Beijing?; Sebastopol, CA.

65. Sima, A.C., de Farias, T.M., Zbinden, E., Anisimova, M., Gil, M., Stockinger, H., Stockinger, K., Robinson-Rechavi, M. and Dessimoz, C. (2019) Enabling Semantic Queries Across Federated Bioinformatics Databases Bioinformatics.

66. Howe, D., Costanzo, M., Fey, P., Gojobori, T., Hannick, L., Hide, W., Hill, D.P., Kania, R., Schaeffer, M., St Pierre, S., et al. (2008) The future of biocuration. Nature, 455, 47–50.

67. International Society for Biocuration (2018) Biocuration: Distilling data into knowledge. PLOS Biol., 16, e2002846.

68. Tang, Y.A., Pichler, K., Füllgrabe, A., Lomax, J., Malone, J., Munoz-Torres, M.C., Vasant, D.V., Williams, E. and Haendel, M. (2019) Ten quick tips for biocuration. PLOS Comput. Biol., 15, e1006906.

69. The SIB Swiss Institute of Bioinformatics’ resources: focus on curated databases (2016) Nucleic Acids Res., 44, D27–D37.

70. Ahi, E.P., Richter, F., Lecaudey, L.A. and Sefc, K.M. (2019) Gene expression profiling suggests differences in molecular mechanisms of fin elongation between cichlid species. Sci. Rep., 9, 9052.

71. Briggs, J.A., Weinreb, C., Wagner, D.E., Megason, S., Peshkin, L., Kirschner, M.W. and Klein, A.M. (2018) The dynamics of gene expression in vertebrate embryogenesis at single-cell resolution. Science, 360, eaar5780.

72. Fushan, A.A., Turanov, A.A., Lee, S.-G., Kim, E.B., Lobanov, A.V., Yim, S.H., Buffenstein, R., Lee, S.-R., Chang, K.-T., Rhee, H., et al. (2015) Gene expression defines natural changes in mammalian lifespan. Aging Cell, 14, 352–365.

73. Holland, P.W., Holland, L.Z., Williams, N.A. and Holland, N.D. (1992) An amphioxus homeobox gene: sequence conservation, spatial expression during development and insights into vertebrate evolution. Dev. Camb. Engl., 116, 653–661.

74. Lam, S.H., Wu, Y.L., Vega, V.B., Miller, L.D., Spitsbergen, J., Tong, Y., Zhan, H., Govindarajan, K.R., Lee, S., Mathavan, S., et al. (2006) Conservation of gene expression signatures between zebrafish and human liver tumors and tumor progression. Nat. Biotechnol., 24, 73–75.

