## Supplementary materials for "The Bgee suite: integrated curated expression atlas and comparative transcriptomics in animals"

### *Supplementary material for* The Bgee suite: integrated curated expression atlas and comparative transcriptomics in animals


SUPPLEMENTARY TABLES 3

Table S1: URLs of resources referred to in the main text 3

SUPPLEMENTARY METHODS 3

Uberon integration 4

Taxon constraints 4

Uberon simplification 4

Cycle removal 4

Improvement of mappings to species-specific ontologies 4

Developmental stage integration 5

Term merging 5

Term ordering 5

part_of relations 6

Insertion into Bgee 6

Curation of expression data 6

Healthy wild-type samples in normal conditions 6

Processing of expression data 7

RNA-Seq data 7

Affymetrix data 7

EST data 8

Integration of *in situ* hybridization data from Model Organism Databases 8

Healthy wild type data 8

Data quality 8

Remapping of annotations 8

Verification of annotations and sex inference 9

Expression ranks and expression scores 10

Rank computation for each data type 10

RNA-Seq 10

Affymetrix 11

In situ hybridization 12
ESTs 12
Integration over all data types and experiments 13
Normalization over all data types and conditions 13
Computation of global weighted mean rank for each gene and condition over all data types 13
GTEx data into Bgee 14
Annotation process 14
GTEx data in Bgee 14
GTEx data in our website 14
GTEx data using BgeeDB R package 15
GTEx filtering criteria 18
Overall summary 18
How did we select GTEx subjects whose samples could be included in Bgee? 19
How did we retrieve status/organ information for subjects from GTEx whose samples could be included in Bgee? 24
How did we check quality for each sample that could be included in Bgee? 24
Criteria of inclusion regarding healthy wild-type data 26
ADDITIONAL RESOURCES 30
BgeeCall R package 30
NCBITaxon ontology 30
Source code 31
Bgee pipeline source code 31
Bgee application source code 31
IQRray source code 31
REFERENCES 31

#### SUPPLEMENTARY TABLES

**Table S1: URLs of resources referred to in the main text**

| Resource | URL |
|---|---|
| GTEx sample exclusion criteria | https://gtexportal.org/home/faq#sampleExclusion |
| Developmental stage ontologies | https://github.com/obophenotype/developmental-stage-ontologies |
| Bgee Anatomical Similarity anotations | https://github.com/BgeeDB/anatomical-similarity-annotations |
| Ensembl 84 used for Bgee 14 | http://mar2016.archive.ensembl.org/ |
| Ensembl Metazoa 30 used for Bgee 14 | http://ensemblgenomes.org/info/release-notes/30 |
| Uberon taxon constraints | https://github.com/obophenotype/uberon/wiki/Taxon-constraints |
| Uberon OWL | http://purl.obolibrary.org/obo/uberon/ext.owl |
| Uberon OWL | http://purl.obolibrary.org/obo/uberon/composite-metazoan.owl |
| Cell Ontology | http://obofoundry.org/ontology/cl.html |
| NCBI Entrez Programming Utilities Help | https://www.ncbi.nlm.nih.gov/books/NBK25501/ |
| NCBI SRA toolkit | https://www.ncbi.nlm.nih.gov/sra/docs/toolkitsoft/ |
| Aspera | https://asperasoft.com/software/clients/ |
| NCBI Authorized Access System | https://www.ncbi.nlm.nih.gov/books/NBK63573/ |
| C. elegans Gross Anatomy Ontology | http://www.obofoundry.org/ontology/wbbt.html |
| C. elegans development ontology | http://www.obofoundry.org/ontology/wbls.html |

#### SUPPLEMENTARY METHODS

The corresponding Bgee pipeline source code and instructions can be found at https://github.com/BgeeDB/bgee_pipeline/tree/master/pipeline/uberon (archived version for Bgee 14 releases: https://github.com/BgeeDB/bgee_pipeline/tree/v14.0/pipeline/uberon). The custom version of Uberon generated for Bgee can be found at https://github.com/BgeeDB/bgee_pipeline/blob/master/generated_files/uberon/ (OBO format: file custom_composite.obo; OWL format: file custom_composite.owl.zip; archived version for Bgee 14 releases: https://github.com/BgeeDB/bgee_pipeline/tree/v14.0/generated_files/uberon).

**Uberon integration**

*Taxon constraints*

In order for automatic reasoners to determine in which taxa an anatomical entity exists, Uberon uses taxon constraints(1)(2). Bgee needs to override some of these taxon constraints. First, because there can be errors in the ontology, which we need to correct. And second, because we need to define taxon constraints for species-specific terms. Indeed, Uberon comes notably in two flavors. One is the ext.owl ontology(3), which is the core version of Uberon, containing the taxon constraint axioms. Another one is the composite-metazoan.owl ontology(4), which merges species ontologies into the structure of Uberon, merging in taxonomic equivalents, relabeling species-specific classes, and also merging in all of the cell ontology(5)(6). Bgee integrates the composite-metazoan version, in order to be able to describe precisely species-specific anatomies, with taxon constraints for generic terms from the core ontology. But the species-specific terms (e.g., ZFA terms for zebrafish) lack taxon constraints. We thus override these taxon constraints to define them for species-specific terms.

*Uberon simplification*

This simplification aims at keeping only the knowledge needed for Bgee, while simplifying it for an easier browsing. We keep in the ontology only some specific relation types (object properties in OWL), and their child relation types (sub-properties): *is_a*, *part_of*, *develops_from*, *transformation_of*. We remove some terms (e.g., CARO:0000006 “material anatomical entity”), while merging their incoming edges with their outgoing edges. We remove some relations that we feel make the ontology more difficult to browse, e.g., UBERON:0000463 “organism substance” *part_of* UBERON:0000468 “multi-cellular organism” (as of Uberon ext release 2019-06-27, this relation is annotated as “"this relationship may be too strong and may be weakened in future"). And we remove some branches we are not interested in, e.g., BFO:0000141 “immaterial anatomical entity”.

*Cycle removal*

We remove cycles between terms in Uberon. All the cycles were caused by a single issue, see https://github.com/obophenotype/uberon/issues/651. See also documentation at https://github.com/BgeeDB/bgee_pipeline/tree/master/pipeline/uberon for a description of the procedure.

*Improvement of mappings to species-specific ontologies*

Many terms from species-specific ontologies, used in MODs providing *in situ* hybridization data, lack a mapping to Uberon; we can thus not integrate them into Bgee without corrections. We correct as many as possible of these missing or incorrect mappings (see https://github.com/obophenotype/uberon/issues/664 and https://github.com/obophenotype/uberon/issues/1288 for a report of the issues discovered for Bgee 14).

**Developmental stage integration**

We describe here our strategy for insertion into the Bgee database. The motivation for design choices are described at https://github.com/obophenotype/developmental-stage-ontologies/blob/master/external/bgee/known_issues.md (archived version for Bgee 14 releases: https://github.com/obophenotype/developmental-stage-ontologies/blob/97fa5b7e176f8c7f72cd5c7115d2fc36f35e8bb3/external/bgee/known_issues.md). The resulting merged developmental stage ontology used in Bgee can be found at https://github.com/obophenotype/developmental-stage-ontologies/blob/master/external/bgee/dev_stage_ontology.obo (archived version for Bgee 14 releases: https://github.com/obophenotype/developmental-stage-ontologies/blob/97fa5b7e176f8c7f72cd5c7115d2fc36f35e8bb3/external/bgee/dev_stage_ontology.obo).

*Term merging*

To merge each species-specific ontology into the structure of Uberon, we add cross-references to associate species-specific terms to Uberon terms they can be merged with. For instance, the term from the human ontology HsapDv:0000010 “gastrula stage” has a cross-reference to the Uberon term UBERON:0000109 “gastrula stage”. HsapDv:0000010 is merged with UBERON:0000109, and all its children are considered as children of UBERON:0000109 (e.g., HsapDv:0000011 “Carnegie stage 06”).

*Term ordering*

Stages are ordered using the relations “preceded_by” or “immediately_preceded_by”. For integration into Bgee, all terms need to be strictly ordered related to each other, as if all terms possessed a relation “immediately_preceded_by” to another term. This is not always the case. For instance, if the relations in the human ontology provided the information that “Carnegie stage 06” is immediately preceded by “Carnegie stage 05”, and that “Carnegie stage 06a” is the first term in temporal ordering among its siblings, and “Carnegie stage 05c” the last; then the Bgee pipeline can infer that “Carnegie stage 06a” is immediately preceded by “Carnegie stage 05c”.

Of note, for some species, stages can be immediately preceded by more than one term. This is the case for instance in worm, where the stage “L4” can be immediately preceded by either “Dauer”, or “L3”. As of Bgee 14, Bgee cannot accommodate such relations, and we conserve only one of them for insertion into the database.

Also, some developmental stages can be asynchronous in an individual, so that the ontology cannot explicitly state what is their temporal ordering. This is for instance the case in fly for the stages embryonic cycle 15 and 16. As of Bgee 14, Bgee needs an absolute ordering of terms for insertion into the database. We thus add “preceded_by” relations to these terms.

*part_of relations*

For insertion into Bgee, a stage must have one parent at most by a *part_of* relation. But the merging of species-specific ontologies into Uberon can cause inconsistencies. For instance, in the ontology for *Drosophila melanogaster* FBdv, FBdv:00005349 “pupal stage”, mapped to UBERON:0000070 “pupal stage”, is a child of FBdv:00007001 “P-stage”. But in Uberon, UBERON:0000070 “pupal stage” is a child of UBERON:0000092 “post-embryonic stage”. The merging of FBdv and Uberon results in having UBERON:0000070 “pupal stage” child of both UBERON:0000092 “post-embryonic stage” and FBdv:00007001 “P-stage”. This is not possible for insertion as of Bgee 14. In this case, we discard the term FBdv:00007001 “P-stage”, and we map all its children to UBERON:0000092 “post-embryonic stage” (FBdv:00005342 “prepupal stage”, FBdv:00005349 “pupal stage”, FBdv:00006011 “pharate adult stage”).

*Insertion into Bgee*

For Bgee 14, we then transform this merged developmental stage ontology into a nested set model (we will likely abandon this approach in future releases, to avoid the issues described above). One of the main difficulties at this step is to order consistently all terms from all species-specific ontologies, related to each other. This cannot be done with a classical sort algorithm, such as merge sort (see (7)). The reason is that some developmental stages in different species cannot be ordered relative to each other. While the classical sorting algorithm considers these stages as equal, it only means that they could not be ordered, not that they truly have the same position in the stage ordering. To overcome this problem, we have implemented a bubble sort algorithm (see (8)), to compare all stages to each other (archived source code for Bgee 14 releases of the sort algorithm: https://github.com/BgeeDB/bgee_apps/blob/Bgee_v14.0/bgee-pipeline/src/main/java/org/bgee/pipeline/ontologycommon/OntologyUtils.java; archived source code for Bgee 14 releases of the sorting algorithm implementation and the generation of the nested set model: https://github.com/BgeeDB/bgee_apps/blob/Bgee_v14.0/bgee-pipeline/src/main/java/org/bgee/pipeline/uberon/UberonDevStage.java).

**Curation of expression data**

*Healthy wild-type samples in normal conditions*

Examples of decisions made concerning conditions which can be considered “normal”.

We have defined whether we could integrate samples used to study the effect of fasting on gene expression: a reasonable fasting time can correspond to conditions that animals encounter in the wild; but an abnormally long fasting time can have adverse effects on the health of the individual, so that it cannot be considered as “healthy” anymore. As a result, we have defined 5-6 hours as being a reasonable fasting time in *Mus musculus*.

Lab strain animals do not exhibit the exact same phenotypes or gene expression patterns than animals harvested in the wild(9), and some lab or domesticated strains are selected to maximize a given phenotype (e.g., DBA/2J mouse model for experimental glaucoma(10)). But they also represent a part of the genetic diversity of the species studied. For this reason, and because lab strain animals represent the vast majority of the data available, we include them into Bgee, while annotating the strain used whenever possible (see below).

**Processing of expression data**

*RNA-Seq data*

We parse Bgee annotations (the files resulting from manual curation of primary databases) in order to retrieve the relevant information about SRA files to download, using the NCBI e-utils(11) (for instance, SRR IDs of runs part of a library). We download data from SRA(12) using the NCBI SRA Toolkit(13) and the Aspera software(14), and then convert them to FASTQ files(15). For GTEx data, we download FASTQ files through the dbGaP(16) Authorized Access System(17) using Aspera. GTF annotation files and genome sequence fasta files from Ensembl(18) and Ensembl metazoa(19) are used to identify sequences of genic regions, exonic regions, and intergenic regions. We also use them to generate indexed transcriptome files for all species in Bgee, using TopHat(20) and Kallisto(21). We use Kallisto to generate pseudo-counts for each transcript, which are then summed for each gene. All further analysis and reporting is done at the gene level as of Bgee 14. TPM and FPKM (or RPKM) values are computed from pseudo-counts, and the results between libraries for each experiment are normalized independently using TMM(22). The source code and documentation for RNA-Seq data integration can be found at https://github.com/BgeeDB/bgee_pipeline/tree/master/pipeline/RNA_Seq (archived version for release Bgee 14.1: https://github.com/BgeeDB/bgee_pipeline/tree/v14.1/pipeline/RNA_Seq).

*Affymetrix data*

We retrieve Affymetrix data preferentially as CEL files, or as MAS5(23) processed files when the CEL files are not available. For each chip type, we retrieve the mapping of probesets to genes from Ensembl. We remove hidden redundancy as explained above. We remove low quality chips by using either the IQRray(24) quality score for CEL file data, or the percentage of genes considered as expressed for MAS5 file data. When CEL files are available for an experiment, we normalize the probeset signal intensities using gcRMA(25). Otherwise, we store the MAS5 flags of expression “present”, “marginal”, “absent”. The source code and documentation for Affymetrix data integration can be found at https://github.com/BgeeDB/bgee_pipeline/tree/master/pipeline/Affymetrix (archived version for release Bgee 14.1: https://github.com/BgeeDB/bgee_pipeline/tree/v14.1/pipeline/Affymetrix).

*EST data*

For ESTs, we retrieve data from UniGene(26) and smiRNAdb(27) (both now retired; Table 1). We retrieve mappings of UniGene data to genomes, for human, mouse, zebrafish and Xenopus, from Ensembl. For fly, as a mapping is not available from Ensembl, we retrieve cDNA information in FASTA format(28) from Ensembl, and use BLAST(29) to map UniGene clusters to genes. In each library, we simply count the number of ESTs mapped to each gene. The source code and documentation for EST data integration can be found at https://github.com/BgeeDB/bgee_pipeline/tree/master/pipeline/ESTs (archived version for release Bgee 14.1: https://github.com/BgeeDB/bgee_pipeline/tree/v14.1/pipeline/ESTs).

**Integration of *in situ* hybridization data from Model Organism Databases**

We retrieve *in situ* hybridization data from the relevant MODs (Table 1). When possible, our pipeline relies on the use of InterMine(30). The source code and documentation for *in situ* hybridization data integration can be found at https://github.com/BgeeDB/bgee_pipeline/tree/master/pipeline/In_situ (archived version for release Bgee 14.1: https://github.com/BgeeDB/bgee_pipeline/tree/v14.1/pipeline/In_situ).

*Healthy wild type data*

For each MOD, we filter out the data from animals with abnormal genetic backgrounds, e.g., animals with transgenes or knockout. We discuss with the developers of these resources to determine how to accurately filter out these data. For instance, for data in *C. elegans*, WormBase has provided us a file containing only healthy wild-type data (Daniela Raciti, personal communication).

*Data quality*

When no data quality information is provided, we assume by default that *in situ* hybridization data are high quality. When the information is provided, we remap the quality levels from each MOD to Bgee quality levels, and discard samples of low quality (for correspondences and filtering criteria, see https://github.com/BgeeDB/bgee_pipeline/tree/master/pipeline/In_situ; archived version for Bgee 14.1, see https://github.com/BgeeDB/bgee_pipeline/tree/v14.1/pipeline/In_situ).

*Remapping of annotations*

Each MOD uses their own controlled vocabulary or ontology to annotate data: EMAPA and EMAPS for GXD in mouse(31); FBbt and FBdv for BDGP and FlyBase in fly(32); ZFA for ZFIN in zebrafish(33); XAO for Xenbase in Xenopus(34, 35); WBbt(36) and WBls(37) for WormBase in worm. For integration into Bgee, we remap these species-specific terms to Uberon, both for the anatomical localizations and the developmental stages. This is achieved by using cross-references or OWL equivalentClass axioms present in Uberon, mapping to these species-specific ontologies. We have also corrected and added many cross-references as part of our integration of Uberon, and of developmental stage ontologies (see above). Other species-specific terms, not merged to Uberon terms, are also present in the composite-metazoan version of Uberon, allowing us to integrate them when needed.

For some MODs, both the anatomy and developmental stage information are entailed in a single term annotation. It is for instance the case in GXD, that uses the EMAPS ontology. The EMAPS ontology maps to anatomical terms from EMAPA, but targets them at a specific developmental stage (see (31) for details). For integration into Bgee, we transform these annotations to retrieve the anatomical localization on the one hand, and the developmental stage on the other hand.

In some other MODs, expression patterns are captured without keeping a stage-specific identity. In WormBase, a gene could be reported as expressed, for instance, in the pharynx at L1 stage and in neurons from embryo to adult; the resulting annotation would be pharynx and neuron, in embryo, larva and adult. The stage specific information is lost. Even when there is only one stage, a gene could be reported as expressed in the pharynx, vulva, and intestine, and, at stage L1, in the nerve ring. Wormbase would capture L1 for nerve ring, but along with the other anatomical parts (Daniela Raciti, personal communication). For this reason, we keep stage information from WomBase only when there is one single anatomical entity annotated for a gene, so that we can be certain that the annotated stages should be associated to it. For other annotations, we map the stage information to the root of our stage ontology, UBERON:0000104 “life cycle”, meaning that no precise information about stage is captured.

The species-specific ontologies used by MODs also differ in their structure, as compared to Uberon, making the mapping task challenging. For instance, the EMAPA ontology makes no distinction between “myocardium” and “cardiac muscles”: “myocardium” is a synonym of the term EMAPA:32688 “cardiac muscle tissues”. EMAPA:32688 is mapped to 3 different terms in Uberon: UBERON:0002349 “myocardium”, UBERON:0001133 “cardiac muscle tissue”, and UBERON:0004493 “cardiac muscle tissue of myocardium”. In this case, Bgee retains the mapping to the most specific term, which is here UBERON:0004493 “cardiac muscle tissue of myocardium”. In other cases, an arbitrary choice has to be made, and only one mapping is kept in the ontology.

##### Verification of annotations and sex inference

Several sanity checks are automatically performed on all annotations, from Bgee and from MODs. We verify that the anatomical entity or developmental stage used in annotation is expected to exist in the species annotated. This checks both our annotations and the Uberon taxon constraints. We verify that the anatomical entity is consistent with the sex annotated, to avoid, for instance, an annotation of “ovary” in “male”. This information is inferred from Uberon, and by using information captured by Bgee about the sexes existing in species (for instance, to know whether hermaphrodite individuals can exist in the species). The inference of the sexes where anatomical entities can exist is reported in our pipeline source code at
https://github.com/BgeeDB/bgee_pipeline/tree/master/generated_files/uberon/uberon_sex_info.tsv
(archived version for release Bgee 14 releases:

https://github.com/BgeeDB/bgee_pipeline/tree/v14.0/generated_files/uberon/uberon_sex_info.tsv). When the sex information is not available for a sample, or we did not capture it at time of annotation, we use the same information to infer the sex automatically. For instance, in species with no hermaphrodite individuals, an “ovary” annotation allows to infer that the sex is “female”. We store whether the sex is inferred, or originally annotated.

**Expression ranks and expression scores**

As of Bgee 14.1, we compute ranks using only data annotated to a condition itself (no propagation), and only considering anatomical entity, developmental stage, and species (i.e., over all sexes and strains indifferently). The rank of a gene in an anatomical entity for a species is the minimum of its individual ranks at each developmental stage with data for it. The pipeline source code and documentation can be found at https://github.com/BgeeDB/bgee_pipeline/tree/master/pipeline/post_processing (archived version for Bgee release 14.1: https://github.com/BgeeDB/bgee_pipeline/tree/v14.1/pipeline/post_processing).

Of note, this pipeline also offers the possibility to compute ranks by pooling all data in a condition itself and its sub-conditions. But we have not yet evaluated and released these ranks, although they would allow to quantify expression levels in many more conditions.

*Rank computation for each data type*

*RNA-Seq*

The Perl script to generate ranks for RNA-Seq data can be found at https://github.com/BgeeDB/bgee_pipeline/blob/master/pipeline/post_processing/ranks_rnaseq.pl (archived version for Bgee release 14.1: https://github.com/BgeeDB/bgee_pipeline/tree/v14.1/pipeline/post_processing/ranks_rnaseq.pl).

To compute ranks for RNA-Seq data, we first identify the set of valid genes that should be considered, and that is always the same for computing ranks in all libraries: the set of all genes that received at least one read over all libraries in Bgee for this species. Then, in each library, we compute fractional ranks of genes, based on their TPM value in this library. For each gene and each condition with RNA-Seq data in the condition itself, we compute a weighted mean of the gene ranks, using all libraries annotated to this condition itself in the species. We weigh the mean by the number of distinct ranks in each library, under the assumption that libraries with a higher number of distinct ranks have a higher power for ranking genes.

To be able to normalize ranks between conditions and data types, we store the maximum fractional rank over all libraries in each condition and species (and not the maximum weighted mean). To be able to compute the global weighted mean rank of a gene over all data types considered in a condition, we store the sum of the number of distinct ranks in each library, for each condition and species. For example, if we have a set of 3 genes to rank, and 2 libraries annotated to the same condition; in library 1, gene 1 has rank 1, gene 2 has rank 2, and gene 3 has rank 3; in library 2, gene 1 has rank 1, gene 2 has rank 2.5, and gene 3 has rank 2.5; the maximum rank in this condition is 3, and the sum of the number of distinct ranks is 3 + 2 = 5.

*Affymetrix*

The Perl script to generate ranks for Affymetrix data as of Bgee 14.1 is available at https://github.com/BgeeDB/bgee_pipeline/blob/master/pipeline/post_processing/ranks_affymetrix.pl (archived version for Bgee release 14.1: https://github.com/BgeeDB/bgee_pipeline/tree/v14.1/pipeline/post_processing/ranks_affymetrix.pl).

To compute ranks for Affymetrix data, we first compute fractional ranks for each gene, for each Affymetrix chip, based on the highest signal intensity from all the probesets mapped to this gene. Then, we normalize ranks between chips annotated to the condition itself in this species. Indeed, different chip types do not have the same probeset design, and thus the same number of genes represented on it. The ranks are then inconsistent between different chip types.

For each chip type, we compute its max rank over all conditions. We normalize ranks of each chip based on the max rank of its corresponding chip type, compared to the max of the max ranks of all chip types used in the same condition. The idea is to correct for the different genomic coverage of different chip types. We do not normalize simply based on the max rank in a given condition, but based on the max rank of the chip types represented in that condition, to not penalize conditions with a lower number of expressed genes, or higher number of ex-aequo genes and lower fractional max ranks. For example, if two chip types are represented in a condition, and the max rank of chipType 1 over all conditions is 1,000, and the max rank of chipType 2 over all conditions is 2,000; the putative rank of a gene represented on chipType 1 could then be shifted, as compared to genes on chipType 2, by a factor of (2000/1000). We thus normalize fractional ranks of genes on chipType 1 by considering the average of their actual rank and their maximal shifted rank: (fractional_rank + fractional_rank * 2000/1000)/2.

We then compute the weighted mean of the normalized ranks per gene and condition, weighting by the number of distinct ranks in each chip, under a similar assumption to RNA-Seq that chips with higher number of distinct ranks have a higher power for ranking genes. Similar to RNA-Seq, to be able to normalize ranks between conditions and data types, we store the max of max ranks of chip types represented in each condition and species. To be able to compute the global weighted mean rank of a gene over all data types considered in a condition, we store the sum of the number of distinct ranks in each chip, for each condition and species.

Of note, for RNA-Seq data, we do not normalize ranks between samples in the same condition before computing the weighted mean, as for Affymetrix data: we use all libraries to produce ranking over always the same set of genes in a given species, so the genomic coverage is always the same, and no normalization is required. The higher power at ranking genes of a library (for instance, thanks to a higher number of mapped reads) is taken into account by weighting the mean by the number of distinct ranks in the library, not by normalizing libraries with lower max ranks; this would penalize conditions with a lower number of expressed genes, and thus with more equally ranked genes, corresponding to genes receiving 0 read.

*In situ hybridization*

Since *in situ* hybridization data are not quantitative, we use an approach based on the assumption that the more often an expression information is reported, the more biologically important this expression is likely to be. The Perl script to generate ranks for *in situ* hybridization data is available at https://github.com/BgeeDB/bgee_pipeline/blob/master/pipeline/post_processing/ranks_in_situ.pl (archived version for Bgee release 14.1: https://github.com/BgeeDB/bgee_pipeline/tree/v14.1/pipeline/post_processing/ranks_in_situ.pl).

To compute ranks for *in situ* hybridization data, we compute a score for each gene in each condition, based on the detection flag of *in situ* evidence (present, absent), and the quality level (high quality, low quality). Each spot is given the following score: [present, high quality], 1; [present, low quality], 0.5; [absent, low quality], -0.5; [absent, high quality], -1. Then we sum these scores for each gene in each condition. We consider all experiments pooled together, because each *in situ* experiment usually studies a very limited number of genes. We compute a dense ranking of the genes in each condition, based on the scores computed. Again, we have considered all experiments pulled together. A fractional ranking is not appropriate, because there are many ex-aequo, leading to an artificially high max rank. This gives too much weight to *in situ* hybridization data when computing a global weighted mean rank over all data types. To be able to normalize ranks between conditions and data types, and to compute the global weighted mean rank of a gene over all data types considered in a condition, we store the max rank in each condition and species.

*ESTs*

While EST data are in principle quantitative, their very low coverage leads us to treat them similarly to *in situ* hybridization data. The Perl script used is available at https://github.com/BgeeDB/bgee_pipeline/blob/master/pipeline/post_processing/ranks_est.pl (archived version for Bgee release 14.1: https://github.com/BgeeDB/bgee_pipeline/tree/v14.1/pipeline/post_processing/ranks_est.pl).

To compute ranks for EST data, we sum for each gene in each condition the count of ESTs mapped to this gene, from all EST libraries in this condition pooled. We compute a dense ranking of the genes in each condition, based on the EST count sums, considering all libraries in the condition all together. A fractional ranking was not appropriate for the same reason as for *in situ* hybridization data. To be able to normalize ranks between conditions and data types, and to compute the global weighted mean rank of a gene over all data types considered in a condition, we store the max rank in each condition and species.

*Integration over all data types and experiments*

*Normalization over all data types and conditions*

Different experiments and techniques might have different power and resolution at ranking genes in a same condition. For instance, some *in situ* hybridization data might allow to rank 5 genes in a condition; while RNA-Seq data in the same condition might allow to rank 30,000 genes, with a max rank of 25,000, and 20,000 distinct ranks. The ranks from different experiments and data types can thus be highly heterogeneous, and difficult to compare as such. To overcome this limitation, we normalize ranks by taking into account the max rank for each data type, condition and species, as compared to the max rank over all data in this species. We thus normalize ranks by considering the average of their actual rank and their maximal shifted rank: (rank + rank * (species_max_rank/data_type_condition_max_rank))/2. The Perl script performing this normalization step is available at https://github.com/BgeeDB/bgee_pipeline/blob/master/pipeline/post_processing/normalize_ranks.pl (archived version for Bgee release 14.1: https://github.com/BgeeDB/bgee_pipeline/tree/v14.1/pipeline/post_processing/normalize_ranks.pl).

We normalize the mean ranks for each gene in each condition and for each data type, based on the max rank for this data type in this condition and species, as compared to the max rank over all conditions and data types in this species. For instance, if the max rank in mouse over all data types and conditions is 30,000, and the dense ranking of a gene in a condition by *in situ* hybridization data is 10, and the max rank by *in situ* hybridization data in this condition in mouse is 30, the dense ranking of the gene will become (10 + 10 * 30,000/30)/2 = 5,005. While this will receive a low weight if there is more informative data (see below), it allows to compute a rank in the absence of other data type in this condition. It also avoids always ranking conditions studied only by *in situ* hybridization data at the top.

*Computation of global weighted mean rank for each gene and condition over all data types*

This computation is done in the Java Bgee API directly, to be able to choose the data types to consider to compute the ranks, based on the user request. The Java source code performing this computation is available at https://github.com/BgeeDB/bgee_apps/blob/master/bgee-dao-sql/src/main/java/org/bgee/model/dao/mysql/expressiondata/MySQLGlobalExpressionCallDAO.java (archived version for Bgee release 14.1: https://github.com/BgeeDB/bgee_apps/blob/Bgee_v14.1/bgee-dao-sql/src/main/java/org/bgee/model/dao/mysql/expressiondata/MySQLGlobalExpressionCallDAO.java).

With the information computed per data type we compute a global weighted mean rank for each gene and condition. All types can be used, or only some. This mean is computed by using the normalized mean ranks or normalized dense ranks for a gene in a condition, from each data type considered. For RNA-Seq and Affymetrix data, if they are considered and data exist in this condition for this gene, the mean rank is weighted by the sum of the number of distinct ranks in this condition and species for the corresponding data type. For *in situ* hybridization and EST data, if they are considered and data exist in this condition for this gene, the dense rank is weighted by the max rank in this condition and species for the corresponding data type.

We transform global weighted mean ranks into expression scores. To compute the expression score of a gene in a condition, we retrieve the max rank over all conditions and data types considered, for this species; and the global weighted mean rank for a gene in a condition, computed over all data types considered and data available, as described above. The expression score is equal to (MaxRank + 1 - meanRank) * (100/maxRank).

**GTEx data into Bgee**

In addition to the continuous growth of transcriptomics datasets, some specific projects produce large amounts of data, generated and accessible in a consistent manner, as, notably, the GTEx project[13]. The GTEx project aims at building a comprehensive resource for tissue-specific gene expression in human. Here we describe how this dataset was integrated into Bgee.

*Annotation process*

We applied a stringent re-annotation process to the GTEx data to retain only healthy tissues and non-contaminated samples, using the information available under restricted-access (see next section *GTEx filtering criteria* for more details). For instance, we rejected all samples for 31% of subjects, deemed globally unhealthy from the pathology report (e.g., drug abuse, diabetes, BMI > 35), as well as specific samples from another 28% of subjects who had local pathologies (e.g., brain from Alzheimer patients). We also rejected samples with contamination from other tissues.

In total, only 50% of samples were kept; these represent a high quality subset of GTEx. All these samples were re-annotated manually to specific Uberon anatomy and aging terms.

*GTEx data in Bgee*

All corresponding RNA-seq were reanalyzed in the Bgee pipeline, consistently with all other healthy RNA-seq from human and other species. These data are being made available both through the website, and through BgeeDB R package (with sensitive information hidden).

*GTEx data in our website*

- Annotations can be retrieved from RNA-Seq human experiments/libraries info. Experiment ID of GTEx is 'SRP012682'.

- Processed expression values, from GTEx only, are available on our FTP (download file).
- Gene expression calls are included into human files
- Each human gene page includes GTEx data if there is any (search a gene here).
- TopAnat analyses can be performs here, which leverage the power of the abundant GTEx data integrated with many smaller datasets to provide biological insight into gene lists.

*GTEx data using BgeeDB R package*

More information and examples can be found on the BgeeDB R package page.

Annotations can be retrieved from RNA-Seq human experiments/libraries information. Experiment ID of GTEx is 'SRP012682'.

```
source("https://bioconductor.org/biocLite.R")

biocLite("BgeeDB")

library(BgeeDB)

bgee <- Bgee$new(species = "Homo_sapiens", dataType = "rna_seq")

myAnnotation <- getAnnotation(bgee)
```

Quantitative expression data and presence calls for GTEx can be loaded.

```
bgee <- Bgee$new(species = "Homo_sapiens", dataType = "rna_seq")

# This step can take a lot of time as all Bgee GTEx data have to be downloaded and
uncompressed.

dataGTEx <- getData(bgee, experimentId = "SRP012682")
```

TopAnat analyses can be performed, which leverage the power of the abundant GTEx data integrated with many smaller datasets to provide biological insight into gene lists.

```
bgee <- Bgee$new(species = "Homo_sapiens")

myTopAnatData <- loadTopAnatData(bgee)

# Retrieve all genes with data in Bgee

allGenes <- unique(row.names(myTopAnatData$gene2anatomy))

# List of genes related to autism and epilepsy from Jabbari 2016
```

```
genesOfInterest <- c("ENSG00000183044", "ENSG00000085563", "ENSG00000006071",
"ENSG00000153086",
    "ENSG00000243989", "ENSG00000156110", "ENSG00000150594", "ENSG00000239900",
    "ENSG00000141385", "ENSG00000038002", "ENSG00000142208", "ENSG00000275199",
    "ENSG00000117020", "ENSG00000163631", "ENSG00000159423", "ENSG00000112294",
    "ENSG00000164904", "ENSG00000033011", "ENSG00000182858", "ENSG00000101901",
    "ENSG00000119523", "ENSG00000214160", "ENSG00000088035", "ENSG00000159063",
    "ENSG00000086848", "ENSG00000242110", "ENSG00000137074", "ENSG00000124198",
    "ENSG00000118520", "ENSG00000198844", "ENSG00000131089", "ENSG00000100299",
    "ENSG00000113273", "ENSG00000004848", "ENSG00000104763", "ENSG00000108381",
    "ENSG00000066279", "ENSG00000138363", "ENSG00000159363", "ENSG00000018625",
    "ENSG00000174437", "ENSG00000182220", "ENSG00000185344", "ENSG00000171953",
    "ENSG00000175054", "ENSG00000158321", "ENSG00000086062", "ENSG00000103507",
    "ENSG00000074582", "ENSG00000176697", "ENSG00000157764", "ENSG00000106009",
    "ENSG00000164061", "ENSG00000169814", "ENSG00000111678", "ENSG00000130921",
    "ENSG00000131943", "ENSG00000197603", "ENSG00000141837", "ENSG00000007402",
    "ENSG00000182389", "ENSG00000198668", "ENSG00000143933", "ENSG00000147044",
    "ENSG00000036828", "ENSG00000110395", "ENSG00000015133", "ENSG00000108691",
    "ENSG00000136861", "ENSG00000008086", "ENSG00000064309", "ENSG00000151849",
    "ENSG00000103995", "ENSG00000173575", "ENSG00000100888", "ENSG00000168539",
    "ENSG000001810", "ENSG00000120903", "ENSG00000101204", "ENSG00000175344",
    "ENSG00000274542", "ENSG00000160716", "ENSG00000114859", "ENSG00000073464",
    "ENSG00000186510", "ENSG00000184908", "ENSG00000188603", "ENSG00000102805",
    "ENSG00000128973", "ENSG00000182372", "ENSG00000278220", "ENSG00000184144",
    "ENSG00000278728", "ENSG00000174469", "ENSG00000166685", "ENSG00000168434",
    "ENSG00000213380", "ENSG00000142173", "ENSG00000173085", "ENSG00000088682",
    "ENSG00000006695", "ENSG00000014919", "ENSG00000047457", "ENSG00000165078",
    "ENSG00000157184", "ENSG00000169372", "ENSG00000147571", "ENSG00000160213",
    "ENSG00000064601", "ENSG00000117984", "ENSG00000174080", "ENSG00000115827",
    "ENSG00000077279", "ENSG00000100150", "ENSG00000181192", "ENSG00000091140",
    "ENSG00000101152", "ENSG00000116675", "ENSG00000116641", "ENSG00000172269",
    "ENSG00000000419", "ENSG00000136908", "ENSG00000179085", "ENSG00000188641",
    "ENSG00000197102", "ENSG00000157540", "ENSG00000101210", "ENSG00000096093",
    "ENSG00000111361", "ENSG00000119718", "ENSG00000070785", "ENSG00000115211",
    "ENSG00000145191", "ENSG00000170370", "ENSG00000133216", "ENSG00000112425",
    "ENSG00000178607", "ENSG00000140374", "ENSG00000105379", "ENSG00000171503",
    "ENSG00000103089", "ENSG00000122591", "ENSG00000145982", "ENSG00000091483",
    "ENSG00000112367", "ENSG00000196924", "ENSG00000162769", "ENSG00000119686",
    "ENSG00000110195", "ENSG00000170345", "ENSG00000125740", "ENSG00000176165",
    "ENSG00000160973", "ENSG00000087086", "ENSG00000179163", "ENSG00000022355",
    "ENSG00000166206", "ENSG00000187730", "ENSG00000113327", "ENSG00000054983",
    "ENSG00000141012", "ENSG00000130005", "ENSG00000171766", "ENSG00000105607",
    "ENSG00000140905", "ENSG00000131095", "ENSG00000170266", "ENSG00000178445",
    "ENSG00000074047", "ENSG00000145888", "ENSG00000109738", "ENSG00000173540",
    "ENSG00000087258", "ENSG00000159921", "ENSG00000111670", "ENSG00000090581",
    "ENSG00000135677", "ENSG00000108433", "ENSG00000171723", "ENSG00000233276",
    "ENSG00000176884", "ENSG00000183454", "ENSG00000273079", "ENSG00000152822",
    "ENSG00000169919", "ENSG00000138796", "ENSG00000170445", "ENSG00000172534",
    "ENSG00000164588", "ENSG00000138622", "ENSG00000213614", "ENSG00000049860",
    "ENSG00000165102", "ENSG00000153187", "ENSG00000158104", "ENSG00000174775",
    "ENSG00000276536", "ENSG00000072506", "ENSG00000114378", "ENSG00000181873",
    "ENSG00000010404", "ENSG00000127415", "ENSG00000134049", "ENSG00000166333",
    "ENSG00000124313", "ENSG00000150995", "ENSG00000120071", "ENSG00000278458",
    "ENSG00000275867", "ENSG00000111262", "ENSG00000169282", "ENSG00000069424",
    "ENSG00000184408", "ENSG00000140015", "ENSG00000151704", "ENSG00000177807",
    "ENSG00000187486", "ENSG00000156113", "ENSG00000281151", "ENSG00000075043",
    "ENSG00000184156", "ENSG00000107147", "ENSG00000243335", "ENSG00000068796",
    "ENSG00000276734", "ENSG00000168280", "ENSG00000185467", "ENSG00000118162",
    "ENSG00000133703", "ENSG00000087299", "ENSG00000196569", "ENSG00000143815",
    "ENSG00000108231", "ENSG00000121897", "ENSG00000138095", "ENSG00000187391",
```

```
"ENSG00000169032", "ENSG00000126934", "ENSG00000109339", "ENSG00000204406",
"ENSG00000090674", "ENSG00000147316", "ENSG00000169057", "ENSG00000081189",
"ENSG00000164073", "ENSG00000168282", "ENSG00000100427", "ENSG00000124615",
"ENSG00000164172", "ENSG00000129255", "ENSG00000178802", "ENSG00000177000",
"ENSG00000198793", "ENSG00000196091", "ENSG00000108784", "ENSG00000072864",
"ENSG00000275911", "ENSG00000125356", "ENSG00000131495", "ENSG00000023228",
"ENSG00000213619", "ENSG00000164258", "ENSG00000115286", "ENSG00000110717",
"ENSG00000167792", "ENSG00000049759", "ENSG00000223957", "ENSG00000204386",
"ENSG00000234343", "ENSG00000228691", "ENSG00000227129", "ENSG00000184494",
"ENSG00000227315", "ENSG00000234846", "ENSG00000196712", "ENSG00000151092",
"ENSG00000187566", "ENSG00000087303", "ENSG00000164190", "ENSG00000156574",
"ENSG00000074181", "ENSG00000141458", "ENSG00000119655", "ENSG00000122585",
"ENSG00000185149", "ENSG00000213281", "ENSG00000179915", "ENSG00000079482",
"ENSG00000116329", "ENSG00000112038", "ENSG00000187848", "ENSG00000135124",
"ENSG00000007168", "ENSG00000125779", "ENSG00000173599", "ENSG00000165194",
"ENSG00000160299", "ENSG00000131828", "ENSG00000148459", "ENSG00000164494",
"ENSG00000127980", "ENSG00000108733", "ENSG00000142655", "ENSG00000121680",
"ENSG00000164751", "ENSG00000215193", "ENSG00000034693", "ENSG00000139197",
"ENSG00000124587", "ENSG00000112357", "ENSG00000102144", "ENSG00000156531",
"ENSG00000092621", "ENSG00000165195", "ENSG00000108474", "ENSG00000165282",
"ENSG00000124155", "ENSG00000060642", "ENSG00000121879", "ENSG00000184381",
"ENSG00000182621", "ENSG00000123560", "ENSG00000140650", "ENSG00000039650",
"ENSG00000108439", "ENSG00000140521", "ENSG00000115138", "ENSG00000131238",
"ENSG00000102103", "ENSG00000139174", "ENSG00000163637", "ENSG00000100033",
"ENSG00000167371", "ENSG00000197746", "ENSG00000185920", "ENSG00000179295",
"ENSG00000172053", "ENSG00000151552", "ENSG00000155961", "ENSG00000132155",
"ENSG00000108557", "ENSG00000146282", "ENSG00000078328", "ENSG00000167281",
"ENSG00000189056", "ENSG00000163933", "ENSG00000155906", "ENSG00000104889",
"ENSG00000136104", "ENSG00000172922", "ENSG00000067836", "ENSG00000151835",
"ENSG00000101347", "ENSG00000138760", "ENSG00000144285", "ENSG00000105711",
"ENSG00000136531", "ENSG00000153253", "ENSG00000183873", "ENSG00000196876",
"ENSG00000169432", "ENSG00000130489", "ENSG00000073578", "ENSG00000178980",
"ENSG00000152217", "ENSG00000127990", "ENSG00000181523", "ENSG00000164690",
"ENSG00000108061", "ENSG00000138083", "ENSG00000064651", "ENSG00000124140",
"ENSG00000119899", "ENSG00000135917", "ENSG00000106688", "ENSG00000110436",
"ENSG00000079215", "ENSG00000102743", "ENSG00000125454", "ENSG00000177542",
"ENSG00000117394", "ENSG00000164414", "ENSG00000117620", "ENSG00000181830",
"ENSG00000076351", "ENSG00000144290", "ENSG00000142319", "ENSG00000276996",
"ENSG00000165970", "ENSG00000130821", "ENSG00000198689", "ENSG00000072501",
"ENSG00000108055", "ENSG00000166311", "ENSG00000102172", "ENSG00000163877",
"ENSG00000115904", "ENSG00000104450", "ENSG00000152583", "ENSG00000166068",
"ENSG00000197694", "ENSG00000102359", "ENSG00000126091", "ENSG00000115525",
"ENSG00000124356", "ENSG00000123473", "ENSG00000136854", "ENSG00000144455",
"ENSG00000139531", "ENSG00000148290", "ENSG00000008056", "ENSG00000197283",
"ENSG00000227460", "ENSG00000102003", "ENSG00000198198", "ENSG00000164458",
"ENSG00000136463", "ENSG00000143374", "ENSG00000054611", "ENSG00000162065",
"ENSG00000145979", "ENSG00000184058", "ENSG00000143178", "ENSG00000196628",
"ENSG00000177426", "ENSG00000175606", "ENSG00000061938", "ENSG00000166340",
"ENSG00000213689", "ENSG00000165699", "ENSG00000103197", "ENSG00000154743",
"ENSG00000274672", "ENSG00000275165", "ENSG00000274796", "ENSG00000274078",
"ENSG00000278712", "ENSG00000273896", "ENSG00000170892", "ENSG00000274129",
"ENSG00000278605", "ENSG00000278622", "ENSG00000182173", "ENSG00000175894",
"ENSG00000104833", "ENSG00000131462", "ENSG00000128159", "ENSG00000198431",
"ENSG00000114062", "ENSG00000104517", "ENSG00000173218", "ENSG00000137411",
"ENSG00000236178", "ENSG00000234032", "ENSG00000206476", "ENSG00000230985",
"ENSG00000223494", "ENSG00000213585", "ENSG00000165637", "ENSG00000197969",
"ENSG00000141252", "ENSG00000196998", "ENSG00000075702", "ENSG00000186153",
"ENSG00000169554", "ENSG00000043355")

# Build the gene vector for the analysis
```

```
geneList <- factor(as.integer(unique(allGenes) %in% genesOfInterest))

names(geneList) <- unique(allGenes)

# Run the test

myTopAnatObject <- topAnat(myTopAnatData, geneList)

resFis <- runTest(myTopAnatObject, algorithm ="elim", statistic ="fisher")

# Format results

tableOver <- makeTable(myTopAnatData, myTopAnatObject, resFis, 0.1)

tableOver
```

*GTEx filtering criteria*

*Overall summary*

1. We selected subjects with biological characteristics matched by Bgee requirements. We created a set of "accepted subjects" by filtering via a subset of variables reported in "phs000424.v6.pht002742.v6.p1.c1.GTEx_Subject_Phenotypes.GRU.txt". We came up with three category tags:
    1.1. "YES": subjects for which we had no evidence for unmatched Bgee requirements
    1.2. "NO": subjects for which we had evidence for large set of unmatched Bgee requirements (eg. systemic disease)
    1.3. "PENDING": subjects for which we had evidence for restricted set of unmatched Bgee requirements (eg. localized disease)
2. We selected samples from "YES" and "PENDING" subjects. The samples were selected from "phs000424.v6.pht002743.v6.p1.c1.GTEx_Sample_Attributes.GRU.txt" file, based on the first part of the GTEx_SampleID (exact string match with GTEx_SubjectID) and created a set of "accepted samples":
    2.1. "YES" subjects: select samples for which we had no evidence for unmatched Bgee requirements, aka select all the samples from this subject
    2.2. "PENDING" subjects: select samples for which we had no evidence for unmatched Bgee requirements
3. We filtered accepted samples:
    3.1. We checked assay type for accepted samples, based on the SMGEBTCHT ['Type of genotype or expression batch'] variable in

"phs000424.v6.pht002743.v6.p1.c1.GTEx_Sample_Attributes.GRU.txt" file. We only accepted TrueSeq.v1(RNA-seq).

3.2. We checked tissue quality for accepted samples, based on free-text SMPTHNTS ['Pathology Notes'] variable in "phs000424.v6.pht002743.v6.p1.c1.GTEx_Sample_Attributes.GRU.txt" file. We predefined a set of quality requirements. We defined a "tissue quality" variable. We came up with two category tags for this variable:

3.2.1. "GOOD": samples for which we had no evidence for unmatched Bgee quality requirements

3.2.2. "BAD": samples for which we had evidence for unmatched Bgee quality requirements

3.3. For all the samples retained so far by our Bgee filters, we then checked the status of the anatomical mapping provided by GTEx (SMUBRID/SMUBRTRM, Uberon ID and Uberon name) by analysing three variables describing tissue location (SMSMPSTE/SMTS/SMTSD) in "phs000424.v6.pht002743.v6.p1.c1.GTEx_Sample_Attributes.GRU.txt" file. We defined a "mapping quality" variable. We came up with two category tags for this variable:

3.3.1. "GOOD": samples for which the Uberon mapping proposed by GTEx was consistent with the tissue location information.

3.3.2. "BAD": samples for which the Uberon mapping proposed by GTEx was not fully consistent with the tissue location information.

4. we then perform a review of Uberon mapping:

4.1. We proposed a new Uberon mapping for filtered samples of "BAD" mapping quality

4.2. We kept as it is the GTEx Uberon mapping for those samples filtered as "GOOD" mapping quality

5. We built the final annotation file, with the samples to be processed into the Bgee pipeline

*How did we select GTEx subjects whose samples could be included in Bgee?*

Only "normal" expression is considered in Bgee (i.e., no treatment, no disease, no gene knock-out...). Since we are dealing with data from human source, it would be too conservative to only keep perfectly healthy subjects, thus we had to find a compromise. We annotated in Bgee only samples from subjects for which we had no evidence for abnormal status (eg. diseases, non-ordinary physiological parameters...).

We excluded from Bgee all samples from subjects suffering from potentially systemic diseases (eg. fungal, bacterial, viral infections), drug/medicine abuse (cocaine, heroin, prescription pills...), abnormally high BMI (above an arbitrarily set threshold of BMI=35), and other "selected" diseases following discussions with annotators (eg. gonorrhea, lupus, sex-transmitted diseases... ); we also excluded subjects for which we had no details on reason of death or if the subject was reported to be ineligible for GTEx. Samples coming from cultured cell lines were also excluded (including EBV-transformed cells). We selectively excluded tissues sampled from unhealthy organs while the subject was otherwise judged to be healthy or affected by a disease not reported to lead to systemic infections in literature: for example, if a subject was reported to be suffering from "ascites", then we excluded liver samples from this subject, but we kept all samples from other organ source for the analyzed subject. We considered as includable in Bgee all samples from individuals which could not be excluded following the given parameters:

We defined a "subject status" variable, harbouring three levels: "YES", "PENDING", O". We assigned one of the levels in variable "status" to each subject provided in GTEx v6.0, depending on the subject's features. We looked up the subject features in the provided file "phs000424.v6.pht002742.v6.p1.c1.GTEx_Subject_Phenotypes.GRU.txt".

In particular, for status assignment of each subject, we were interested in the following columns:

- BMI : Autocalculated field of BMI: general indicator of the body fat an individual is carrying based upon the ratio of weight to height. We excluded from Bgee all samples from subjects having a BMI value greater than 35.

- INCEXC : A verification field of whether the Donor has met the overall eligibility criteria for GTEx collection based on answers to eligibility questions. We excluded from Bgee all samples from subjects that were reported "false" for this field.

- DTHFUCOD / DTHCOD: First Underlying Cause Of Death / Immediate Cause of Death. We excluded from Bgee all samples from subjects that were reported to died from: drug/medicine abuse/intoxication, cancer, end-stage diseases (eg. kidney, liver...)

- DTHHRDY: Death Classification: 4-point Hardy Scale. We excluded from Bgee all samples from subjects that were tagged with a Hardy Scale level of 4 (aka: Slow death Death after a long illness, with a terminal phase longer than 1 day -commonly cancer or chronic pulmonary disease-; deaths that are not unexpected) only if DTHFUCOD / DTHCOD and DTHHRDY are consistent (eg: a subject had pneumonia and died of pneumonia, DTHHRDY 4).

- LBHBCABM / LBHBCABT / LBHBSAB / LBHBSAG: Hepatitis B virus infection serology analysis. We excluded from Bgee all samples from subjects that were tagged as "positive" (=1) in at least one of these columns.

- LBHCV1NT / LBHBHCVAB: Flaviviridae infection serology analysis. We excluded from Bgee all samples from subjects that were tagged as "positive" (=1) in at least one of these columns.

- LBHBHCVAB / LBHIV1NT / LBHIVAB / LBHIVO: HIV serology analysis. We excluded from Bgee all samples from subjects that were tagged as "positive" (=1) in at least one of these columns.

- LBPRRVDRL / LBRPR: Syphilis or syphilis agent serology analysis. We excluded from Bgee all samples from subjects that were tagged as "positive" (=1) in at least one of these columns.

- MHALS: Amytrophis Lateral Sclerosis. We excluded from Bgee all samples from subjects that were tagged as "positive" (=1) in this column.

- MHALZDMT / MHALZHMR: Alzheimer. We excluded from Bgee brain samples coming from subjects that were tagged as "positive" (=1) in at least one of these columns.

- MHARTHTS: Arthritis. We excluded from Bgee all samples from subjects that were tagged as "positive" (=1) in this column.

- MHASCITES: Ascites. We excluded from Bgee liver samples coming from subjects that were tagged as "positive" (=1) in this column.

- MHBCTINF: Bacterial infection. We excluded from Bgee all samples from subjects that were tagged as "positive" (=1) in this column.

- MHCANCER5 / MHCANCERC / MHCANCERNM: Cancer detection or cancer history. We excluded from Bgee all samples from subjects that were tagged as "positive" (=1) in this column.

- MHCLLULTS: Cellulites. We excluded from Bgee skin samples coming from subjects that were tagged as "positive" (=1) in this column.

- MHCOCAINE5: Cocaine use/abuse in the 5 years before death. We excluded from Bgee all samples from subjects that were tagged as "positive" (=1) in this column.

- MHCOPD: Chronic low-respiratory disease. We excluded from Bgee lung samples coming from subjects that were tagged as "positive" (=1) in this column.

- MHCOUGHU: Chronic respiratory disease. We excluded from Bgee lung samples coming from subjects that were tagged as "positive" (=1) in this column.

- MHCVD: Cerebrovascular disease/stroke/encephalitis. We excluded from Bgee brain samples coming from subjects that were tagged as "positive" (=1) in this column.

- MHDLYSIS: Dialysis. We excluded from Bgee all samples from subjects that were tagged as "positive" (=1) in this column.

- MHDMNTIA: Dementia. We excluded from Bgee brain samples coming from subjects that were tagged as "positive" (=1) in this column.

- MHENCEPHA: Active encephalitis. We excluded from Bgee brain samples coming from subjects that were tagged as "positive" (=1) in this column.

- MHFNGINF: Fungal infections. We excluded from Bgee all samples from subjects that were tagged as "positive" (=1) in this column.

- MHGNRR12M: Gonhorrea. We excluded from Bgee all samples from subjects that were tagged as "positive" (=1) in this column.

- MHHEPBCT: Hepatitis B infection. We excluded from Bgee all samples from subjects that were tagged as "positive" (=1) in this column.

- MHHEPCCT: We excluded from Bgee all samples from subjects that were tagged as "positive" (=1) in this column.

- MHHEROIN: Heroin users. We excluded from Bgee all samples from subjects that were tagged as "positive" (=1) in this column.

- MHHMPHLIA / MHHMPHLIAB: Hemophilia. We excluded from Bgee blood sampless from subjects that were tagged as "positive" (=1) in this column.

- MHHRTDIS: Ischemic heart disease. We excluded from Bgee heart samples coming from subjects that were tagged as "positive" (=1) in this column.

- MHHRTDISB: Heart disease history. We excluded from Bgee heart samples coming from subjects that were tagged as "positive" (=1) in this column.

- MHIVDRG5: Intravenous drug abuse. We excluded from Bgee all samples from subjects that were tagged as "positive" (=1) in this column.

- MHLUPUS: Lupus.  We excluded from Bgee all samples from subjects that were tagged as "positive" (=1) in this column.

- MHLVRDIS: Liver chronic disease. We excluded from Bgee liver samples coming from subjects that were tagged as "positive" (=1) in this column.

- MHMENINA: Meningitis.  We excluded from Bgee brain samples coming from subjects that were tagged as "positive" (=1) in this column.

- MHOPPINF: Opportunistic infections.  We excluded from Bgee all samples from subjects that were tagged as "positive" (=1) in this column.

- MHOSTMYLTS: Osteomyelitis. We excluded from Bgee bone samples coming from subjects that were tagged as "positive" (=1) in this column.

- MHPLLABS: Prescription pills abuse. We excluded from Bgee all samples from subjects that were tagged as "positive" (=1) in this column.

- MHPNMIAB / MHPNMNIA: Penumonia. We excluded from Bgee lung samples coming from subjects that were tagged as "positive" (=1) in this column.

- MHPRKNSN: Parkinson. We excluded from Bgee all samples from subjects that were tagged as "positive" (=1) in this column.

- MHPSBLDCLT: Blood culture shown positive to bacteria. We excluded from Bgee all samples from subjects that were tagged as "positive" (=1) in this column.

- MHRA: Arthritis rheumatoides.  We excluded from Bgee all samples from subjects that were tagged as "positive" (=1) in this column.

- MHRNLFLR: Renal failure. We excluded from Bgee kindney samples coming from subjects that were tagged as "positive" (=1) in this column.

- MHSCHZ: Schizophrenia. We excluded from Bgee brain samples coming from subjects that were tagged as "positive" (=1) in this column.

- MHSCLRDRM: Scleroderma, aka systemic sclerosis. We excluded from Bgee all samples from subjects that were tagged as "positive" (=1) in this column.

- MHSDRGABS: Signs of drug abuse. We excluded from Bgee all samples from subjects that were tagged as "positive" (=1) in this column.

- MHSEPSIS: Sepsis. We excluded from Bgee all samples from subjects that were tagged as "positive" (=1) in this column.

- MHSTD: Sex-transmitted diseases. We excluded from Bgee all samples from subjects that were tagged as "positive" (=1) in this column.

- MHSTRDLT: Long-term steroid use. We excluded from Bgee all samples from subjects that were tagged as "positive" (=1) in this column.

- MHSUBABSA / MHSUBABSB: Drug use for non-medical reasons. We excluded from Bgee all samples from subjects that were tagged as "positive" (=1) in this column.

- MHT1D: Diabete type I "genetic".  We excluded from Bgee all samples from subjects that were tagged as "positive" (=1) in this column.

- MHTBHX: Tubercolosis. We excluded from Bgee all samples from subjects that were tagged as "positive" (=1) in this column.

In addition, we decided to:

- exclude brain samples from subjects who died from CerebroVascular Accident (CVA) and whose death was reported on Hardy Scale 3 or 4

According to information reported in the described columns: if samples from a given subject had to be completely excluded from Bgee, then the subject was assigned status "NO"; if some but not all the samples from a given subject had to be excluded from Bgee, then the subject was assigned status

"PENDING"; if samples from a subject had no reason to be excluded from Bgee, then the subject was assigned status "YES".

*How did we retrieve status/organ information for subjects from GTEx whose samples could be included in Bgee?*

For the final annotation file, we needed to retrieve information such as organ source or the subject's attributes such as age, sex, ethnicity...etc.

Useful columns reported in file "phs000424.v6.pht002742.v6.p1.c1.GTEx_Subject_Phenotypes.GRU.txt" were the following:

- SUBJID : Subject ID, GTEx Public Donor ID

- GENDER : The Donor's Identification of gender based upon self-report, family/next of kin, or medical record abstraction. Note: values from this variable were used to obtain the "Sex" value for the final RNAseqLibrary.tsv file.

- AGE : Elapsed time since birth in years. Note: values from this variable were used to obtain the "InfoStage" value for the final RNAseqLibrary.tsv file.

- RACE : Report of Donor's race (geographically based) as reported by Donor, family/next of kin, or medical record abstraction. Note: values from this variable were used to obtain the "Strain" value for the final RNAseqLibrary.tsv file.

Useful columns reported in file "phs000424.v6.pht002743.v6.p1.c1.GTEx_Sample_Attributes.GRU.txt" were the following:

- SAMPID: Sample ID, GTEx Public Sample ID

- SMSMPSTE: Tissue Location

- SMTS: Tissue Type, area from which the tissue sample was taken.

- SMTSD: Tissue Type, more specific detail of tissue type

note: values from these three variables were merged together to obtain the "InfoOrgan" value for the final RNAseqLibrary.tsv file.

*How did we check quality for each sample that could be included in Bgee?*

Only good quality samples have to be included in Bgee. Quality level for each sample was assessed for both biological quality, inferred from microscopy sample subset observations by MDs, and annotation mapping quality.

We defined a "Tissue quality" variable, harbouring two levels: "GOOD" or "BAD", to define biological quality of each sample. We defined a "Mapping quality" variable, harbouring two levels: "GOOD", "BAD", to define annotation mapping quality of each sample.

We assigned one of the levels of both "Tissue quality" and "Mapping quality" variables to each sample to be included in Bgee, depending on the sample's features. We looked up the sample features in the provided file "phs000424.v6.pht002743.v6.p1.c1.GTEx_Sample_Attributes.GRU.txt".

We selected only RNA-seq data from GTEx, because they're the ones that the current Bgee pipeline is able to deal with. To select only samples processed with RNA-seq, we selected:

- SMGEBTCHT (Type of genotype or expression batch) = "TrueSeq.v1"

In particular, for "Tissue quality", we were interested in columns:

- SMSTYP: Indicates whether sample is a tumor or a normal. We excluded from Bgee all tumoral samples.

- SMPTHNTS: Pathologist notes, free-text field. We considered as "BAD" samples that contained more than 10% of foreign tissue, as well as samples displaying diseases or early disease stage.

For "Mapping quality", we were interested in columns:

- SMSMPSTE: The location on the organ or tissue from which the specimen was collected.

- SMTS: The type of Tissue that was collected

- SMTSC: Any comments the prosector wishes to make concerning this case with respect to tissue collection (this field contains for example information about the organ sub-anatomy, eg. left/rigt)

- SMTSD: The type of Tissue that was collected on a Donor by Donor basis

- SMUBRID/SMUBRTRM: Uberon term and Uberon ID proposed by annotators.

We considered that a sample had "GOOD" mapping quality if the columns SMSMPSTE/SMTS/SMTSD were consistent with the suggested SMUBRTRM value and if the eventual detail reported in SMTSC was of no biological relevance. Otherwise, we considered that a sample had "BAD" mapping quality if the annotation was considered not precise enough, or if SMUBRTRM was inconsitent with what reported in SMSMPSTE/SMTS/SMTSD, or if there was a biological difference, with possible precision found in Uberon, that was not accurately reported. For all samples with "BAD" mapping quality, we proposed a new Uberon term (and identifier) only if the sample was reported to be of "GOOD" quality.

**Criteria of inclusion regarding healthy wild-type data**

| **Factor** | **Include into Bgee?** | **Comment** | **Example** |
|---|---|---|---|
| BMI (Body Mass Index) from 18.5 to 25 ~~(or less than 30)~~ (or less than 35) | Yes | In the normal range or overweight (overweight is common), see https://en.wikipedia.org/wiki/Body_mass_index to exclude other ranges (Sept, 2019, Update: see comment here above about GTEx samples, we have to consider BMI value is continuously rising inside human population). Actually, weight difference between individuals may be part of the natural variability | SRP046752 |
| Cell lines (3T3-L1, Hela, MCF-7) and cell cultures | No | | |
| Fasted animals | Yes | If the fastening time is reasonable: a mouse fasting duration of 5–6 h might offer a better comparison to humans overnight(16–18 h), see PMID:24025567 | E-GEOD-7137 |
| Dark/light circadian rhythms and temperature variation | yes/no | Depends on the animal, if reasonable for its physiology, yes as in the example | GSE23528 |
| Low or high fat diet for short time | Yes | If the diet time is short (3 days), can be considered as part of the wild life variability for animals | E-GEOD-8524 |

| | | | |
|---|---|---|---|
| Mammary glands from virgin, pregnant and lactating females | Yes | From all types of females | E-TABM-199 |
| Oocytes at different stages of maturation | Yes | Without including the info on the stage (except for Drosophila where the different maturation stages are present in the ontology) | E-GEOD-3351 |
| Placenta and extraembryonic components during development | Yes | Be careful whether to put it in the adult or the embryo! | E-GEOD-7674 |
| Injury | No | | |
| Animals selected for their behaviour (e.g. fear) | Yes | Part of the natural variability | E-GEOD-4035 |
| Animals from different strains (e.g. C57BL/6, BALB/c...) | Yes | Part of the natural variability | |
| Intestinal germ free animals | No | Normal animals have an intestinal flora | E-GEOD-5156 |

| | | | |
|---|---|---|---|
| Removal of the eye (monocular enucleation) or cochlea on one side; the eye, visual cortex or cochlea on the other side were analysed | No | | E-GEOD-4265 |
| Cell types (T-cells, stem cells...) | Yes | Should be included only if enough precision in the ontologies (e.g. T-cell in zebrafish). If not enough precision in the ontology, store the experiment ID in the "not_included_for_now" file | |
| Polysomal RNA only hybridized | No | Method that pellets the polyribosomes while leaving the mono and non-polysomal mRNA fractions in the supernatant. | E-GEOD-3962 |
| Anesthesia | No/Yes | Similar to drug treatment (but sometimes anesthesia is simply not described, and anyway has to be accepted for human tissue samples) | |
| Human post-mortem tissues | Yes | Mainly the simple way to get human tissues | |
| Light impulse to stress the animals | Yes | Stress is probably usual for lab animals | |
| Killed by cervical dislocation or decapitation | Yes | Common for Mouse | |
| Killed by inhalants (CO2) | Yes | Common for Mouse | |

| | | | |
|---|---|---|---|
| Killed by exsanguination under CO2 anesthesia | No | Used for mouse lung retrieval | |
| Killed by intravenous anesthetic | No | Method of killing is sometimes simply not described | |
| Killed by intracardiac or intraperitoneal injection | No | Method of killing is sometimes simply not described | |
| Mock-treated (plasmid, surgery...) | No/Yes | Typical 'control' that is not normal condition, to check. Can be acceptable when this is for example 'mock inoculated in a similar manner with Minimal Essential Medium' (SRP061418), but rejected when involving experimental surgery (SRP039511) | SRP061418, SRP039511 |
| Normal adjacent tissues from tumor | No/Yes | Sometimes difficult to get the information, be careful if both tumor and normal samples come from the same patient (paired samples). Also sometimes we consider the data because no other data are available for these tissues (described in comments) | |
| Treatment with Alkaline Hypochlorite Solution ("Bleaching") of C.elegans worm: larva are collected | Yes | Use for synchronizing C. elegans culture, see here | |

#### ADDITIONAL RESOURCES

In order to serve the development of Bgee, we have contributed to the development of additional resources, listed here: i) a Bioconductor(38) R(39) package allowing to produce present/absent expression calls from RNA-Seq data; and ii) an ontology capturing the information present in the NCBI Taxonomy database(40). In addition, all source code produced for Bgee is publicly available.

##### BgeeCall R package

The BgeeCall R package allows to reproduce a specific part of the Bgee pipeline: the generation of present/absent expression calls from RNA-Seq data (see *Material and methods*, *Calls of presence/absence of expression*, *Calls produced from RNA-Seq data*). Our method avoids the use of a fixed threshold to consider a gene as expressed (such as 1 RPKM, or 2 TPM). It is based on the estimation of background experimental and transcriptional noise from expression level of intergenic regions (Julien Roux, Marta Rosikiewicz, Julien Wollbrett, Marc Robinson-Rechavi, Frederic B. Bastian; in preparation). The BgeeCall R package allows to generate present/absent calls for any RNA-Seq library, as long as the species studied is present in Bgee. It is also possible to use it for other species, if a set of validated intergenic regions is provided (see the Zenodo 'bgee_intergenic' community(41) for more details). The BgeeCall Bioconductor page can be accessed at https://bioconductor.org/packages/BgeeCall/.

##### NCBITaxon ontology

In order to compare expression patterns between species, and to identify homologous anatomical entities to do so, we need to be able to retrieve the common ancestor of any set of species in Bgee. Moreover, to define taxon constraints in a multi-species ontology, a taxonomy ontology is needed (see, e.g., (42) and (43)). To this aim, the NCBI taxonomy database(40) has been translated into an ontology (and more specifically, a tree, where each term has only one parent, except the root taxon "cellular organisms", which has no parent). Each term in the ontology represents a taxon; and each taxon is linked to its direct ancestral taxon by the OBO *is_a* relation (equivalent to *SubClassOf* in OWL). The ontology is part of the OBOFoundry(44), and is available at http://obofoundry.org/ontology/ncbitaxon.html.

But in order to compute taxon constraints in Uberon (see *Material and methods*, *Uberon integration*, *Taxon constraints*), we needed to use a version of the taxon ontology that includes disjoint classes axioms between sibling taxa(45). These axioms are not included in the public version of the ontology. We have thus produced a custom version, including these axioms, for the species present in Bgee. This version can be found at https://github.com/BgeeDB/bgee_pipeline/blob/master/generated_files/species/bgee_ncbitaxon.owl (archived version for Bgee 14 releases: https://github.com/BgeeDB/bgee_pipeline/blob/v14.0/generated_files/species/bgee_ncbitaxon.owl).

#### Source code

##### *Bgee pipeline source code*

The source code of the entire Bgee pipeline is available at https://github.com/BgeeDB/bgee_pipeline/. The pipeline is decomposed in different steps, focused on a specific part of the pipeline (e.g., genome integration, RNA-Seq analyses). For each step, we have created a Makefile(46), allowing to list all the computations necessary, and the dependencies between the computations. And for each step, a README file documents it directly in the step folder.

##### *Bgee application source code*

The source code of the Bgee application is available at https://github.com/BgeeDB/bgee_apps. The Bgee application is developed in Java(47). It is structured in different modules: *bgee-dao-api* defines the interfaces to access to the data source; *bgee-dao-sql* is an implementation of *bgee-dao-api* to access the Bgee MySQL database; *bgee-core* is the business layer of the application, or model layer in a model-view-controller architecture(48), making use of *bgee-dao-api*; *bgee-pipeline* contains the Java components used in the Bgee pipeline, making use of *bgee-core* and *bgee-dao-sql*; *bgee-webapp* contains the code to build the Bgee website, using the *bgee-core* module, representing the view and controller layers in a model-view-controller architecture.

##### *IQRray source code*

To perform quality controls of Affymetrix data where raw CEL files are available, we have developed the IQRray(24) method. IQRray outperforms other methods in identification of poor quality arrays in datasets composed of arrays from many independent experiments. IQRray is implemented in R, and the source code is available at https://github.com/BgeeDB/IQRray.

#### REFERENCES


http://heim.ifi.uio.no/~trygver/themes/mvc/mvc-index.html

1. Deegan (née Clark),J.I., Dimmer,E.C. and Mungall,C.J. (2010) Formalization of taxon-based constraints to detect inconsistencies in annotation and ontology development. *BMC Bioinformatics*, **11**, 530.
2. Chris Mungall,J.D. Taxon constraints in Uberon.
3. Uberon ext.owl.
4. Uberon composite-metazoan.owl.
5. Meehan,T.F., Masci,A.M., Abdulla,A., Cowell,L.G., Blake,J.A., Mungall,C.J. and Diehl,A.D. (2011) Logical Development of the Cell Ontology. *BMC Bioinformatics*, **12**, 6.
6. Cell Ontology.
7. Bongiovanni,G., Bovet,D.P. and Di Battista,G. eds. (1997) Algorithms and complexity: third Italian conference, CIAC '97, Rome, Italy, March 12-14, 1997: proceedings Springer, Berlin ; New York.
8. Astrachan,O. (2003) Bubble sort: an archaeological algorithmic analysis. *ACM SIGCSE Bull.*, **35**, 1.
9. Yoshiki,A. and Moriwaki,K. (2006) Mouse phenome research: implications of genetic background. *ILAR J.*, **47**, 94–102.
10. Turner,A.J., Vander Wall,R., Gupta,V., Klistorner,A. and Graham,S.L. (2017) DBA/2J mouse model for experimental glaucoma: pitfalls and problems. *Clin. Experiment. Ophthalmol.*, **45**, 911–922.
11. Entrez Programming Utilities Help (2010).

12. Kodama,Y., on behalf of the International Nucleotide Sequence Database Collaboration, Shumway,M., on behalf of the International Nucleotide Sequence Database Collaboration, Leinonen,R. and on behalf of the International Nucleotide Sequence Database Collaboration (2011) The sequence read archive: explosive growth of sequencing data. *Nucleic Acids Res.*, **40**, D54–D56.
13. SRA Toolkit download.
14. Aspera software.
15. Cock,P.J.A., Fields,C.J., Goto,N., Heuer,M.L. and Rice,P.M. (2009) The Sanger FASTQ file format for sequences with quality scores, and the Solexa/Illumina FASTQ variants. *Nucleic Acids Res.*, **38**, 1767–1771.
16. Tryka,K.A., Hao,L., Sturcke,A., Jin,Y., Wang,Z.Y., Ziyabari,L., Lee,M., Popova,N., Sharopova,N., Kimura,M., *et al.* (2013) NCBI's Database of Genotypes and Phenotypes: dbGaP. *Nucleic Acids Res.*, **42**, D975–D979.
17. GaP FAQ Archive - Authorized Access System Login (2009).
18. Yates,A.D., Achuthan,P., Akanni,W., Allen,J., Allen,J., Alvarez-Jarreta,J., Amode,M.R., Armean,I.M., Azov,A.G., Bennett,R., *et al.* (2019) Ensembl 2020. *Nucleic Acids Res.*, 10.1093/nar/gkz966.
19. Howe,K.L., Contreras-Moreira,B., De Silva,N., Maslen,G., Akanni,W., Allen,J., Alvarez-Jarreta,J., Barba,M., Bolser,D.M., Cambell,L., *et al.* (2019) Ensembl Genomes 2020-enabling non-vertebrate genomic research. *Nucleic Acids Res.*, 10.1093/nar/gkz890.
20. Kim,D., Pertea,G., Trapnell,C., Pimentel,H., Kelley,R. and Salzberg,S.L. (2013) TopHat2: accurate alignment of transcriptomes in the presence of insertions, deletions and gene fusions. *Genome Biol.*, **14**, R36.
21. Bray,N.L., Pimentel,H., Melsted,P. and Pachter,L. (2016) Near-optimal probabilistic RNA-seq quantification. *Nat. Biotechnol.*, **34**, 525.
22. Robinson,M.D. and Oshlack,A. (2010) A scaling normalization method for differential expression analysis of RNA-seq data. *Genome Biol.*, **11**, R25.
23. Hubbell,E., Liu,W.-M. and Mei,R. (2002) Robust estimators for expression analysis. *Bioinformatics*, **18**, 1585–1592.
24. Rosikiewicz,M. and Robinson-Rechavi,M. (2016) IQRray, a new method for Affymetrix microarray quality control, and the homologous organ conservation score, a new benchmark method for quality control metrics. *Bioinformatics*, **32**, 2565–2565.
25. Wu,Z., Irizarry,R.A., Gentleman,R., Martinez-Murillo,F. and Spencer,F. (2004) A Model-Based Background Adjustment for Oligonucleotide Expression Arrays. *J. Am. Stat. Assoc.*, **99**, 909–917.
26. Pontius,J.U., Wagner,L. and Schuler,G.D. (2004) UniGene: A unified view of the transcriptome. In: The NCBI Handbook National Center for Biotechnology Information.
27. Landgraf,P., Rusu,M., Sheridan,R., Sewer,A., Iovino,N., Aravin,A., Pfeffer,S., Rice,A., Kamphorst,A.O., Landthaler,M., *et al.* (2007) A Mammalian microRNA Expression Atlas Based on Small RNA Library Sequencing. *Cell*, **129**, 1401–1414.
28. Lipman,D. and Pearson,W. (1985) Rapid and sensitive protein similarity searches. *Science*, **227**, 1435–1441.
29. Altschul,S.F., Gish,W., Miller,W., Myers,E.W. and Lipman,D.J. (1990) Basic local alignment search tool. *J. Mol. Biol.*, **215**, 403–410.
30. Kalderimis,A., Lyne,R., Butano,D., Contrino,S., Lyne,M., Heimbach,J., Hu,F., Smith,R., Štěpán,R., Sullivan,J., *et al.* (2014) InterMine: extensive web services for modern biology. *Nucleic Acids Res.*, **42**, W468–W472.
31. Hayamizu,T.F., Baldock,R.A. and Ringwald,M. (2015) Mouse anatomy ontologies: enhancements and tools for exploring and integrating biomedical data. *Mamm. Genome*, **26**, 422–430.
32. Costa,M., Reeve,S., Grumbling,G. and Osumi-Sutherland,D. (2013) The Drosophila anatomy ontology. *J. Biomed. Semant.*, **4**, 32.
33. Van Slyke,C.E., Bradford,Y.M., Westerfield,M. and Haendel,M.A. (2014) The zebrafish anatomy and stage ontologies: representing the anatomy and development of Danio rerio. *J. Biomed. Semant.*, **5**, 12.
34. Segerdell,E., Bowes,J.B., Pollet,N. and Vize,P.D. (2008) An ontology for Xenopus anatomy and development. *BMC Dev. Biol.*, **8**, 92.
35. Segerdell,E., Ponferrada,V.G., James-Zorn,C., Burns,K.A., Fortriede,J.D., Dahdul,W.M., Vize,P.D. and Zorn,A.M. (2013) Enhanced XAO: the ontology of Xenopus anatomy and development underpins more accurate annotation of gene expression and queries on Xenbase. *J. Biomed. Semant.*, **4**, 31.

36. C. elegans Gross Anatomy Ontology.
37. C. elegans Development Ontology.
38. Huber,W., Carey,V.J., Gentleman,R., Anders,S., Carlson,M., Carvalho,B.S., Bravo,H.C., Davis,S., Gatto,L., Girke,T., *et al.* (2015) Orchestrating high-throughput genomic analysis with Bioconductor. *Nat. Methods*, **12**, 115–121.
39. R Core Team (2018) R: A Language and Environment for Statistical Computing R Foundation for Statistical Computing, Vienna, Austria.
40. Federhen,S. (2011) The NCBI Taxonomy database. *Nucleic Acids Res.*, **40**, D136–D143.
41. Zenodo Bgee intergenic community.
42. Kuśnierczyk,W. (2008) Taxonomy-based partitioning of the Gene Ontology. *J. Biomed. Inform.*, **41**, 282–292.
43. Deegan (née Clark),J.I., Dimmer,E.C. and Mungall,C.J. (2010) Formalization of taxon-based constraints to detect inconsistencies in annotation and ontology development. *BMC Bioinformatics*, **11**, 530.
44. Smith,B., Ashburner,M., Rosse,C., Bard,J., Bug,W., Ceusters,W., Goldberg,L.J., Eilbeck,K., Ireland,A., Mungall,C.J., *et al.* (2007) The OBO Foundry: coordinated evolution of ontologies to support biomedical data integration. *Nat. Biotechnol.*, **25**, 1251.
45. Chris Mungall (2012) Taxon constraints in OWL. *Monkeying OWL*.
46. Feldman,S.I. (1979) Make—A program for maintaining computer programs. *Softw. Pract. Exp.*, **9**, 255–265.
47. Gosling,J., Joy,B., Steele,G.L., Bracha,G. and Buckley,A. (2014) The Java Language Specification, Java SE 8 Edition. 1st Addison-Wesley Professional.
48. Reenskaug,T. (1979) A note on DynaBook requirements. *Xerox PARC*.
